# TRIM21 is a molecular rheostat for influenza A virus replication

**DOI:** 10.64898/2026.03.16.711963

**Authors:** Qi Wen Teo, Huibin Lv, Tossapol Pholcharee, Xin Chen, Vidia Ramadin, Kevin J. Mao, Jessica J. Huang, Joel Rivera-Cardona, Jie Zhu, Yang Wei Huan, Evan K. Shao, Hein M. Tun, Christopher B. Brooke, Lilliana Radoshevich, Sumana Sanyal, Nicholas C. Wu

## Abstract

TRIM21 is a multifunctional E3 ubiquitin ligase and intracellular antibody receptor, yet its role during viral infection remains unclear, with reports describing both antiviral and proviral activities. Here, we show that TRIM21 regulates influenza infection in an expression-dependent manner by functioning as a molecular rheostat rather than a binary restriction factor. This graded activity of TRIM21, which leads to both suppression and promotion of influenza replication, couples linkage-specific ubiquitination of viral nucleoprotein with modulation of innate immune signaling. Additionally, loss of TRIM21 unmasks a compensatory antiviral program centered on PRKDC, which is a ubiquitination target of TRIM21. This positions PRKDC as a latent restriction factor selectively engaged when primary TRIM21 control is lost. Together, these findings reveal a hierarchical and plastic antiviral network in which TRIM21 sets an adjustable threshold for host defense while restraining secondary restriction pathways. This framework highlights the sophisticated layers of regulation of the host ubiquitin-mediated antiviral immunity.

## INTRODUCTION

Influenza A virus (IAV) remains a persistent global health threat, due to its rapid evolution and capacity for cross-species transmission. The emergence of highly pathogenic H5N1 with enhanced ability to cross species barriers^1^, together with the identification of a single mutation in bovine H5N1 hemagglutinin (HA) that shifts receptor specificity toward human receptor^2^, underscores the continual emergence of viral variants with altered pathogenic potential. These evolving threats highlight the importance of understanding the host defense mechanisms that constrain viral replication while limiting immunopathology. Such control is achieved through multilayered regulatory networks rather than single dominant restriction factors, with the ubiquitin system serving as a central coordinator of innate immune signaling, inflammatory responses, and protein quality control^3–5^.

Ubiquitination is a reversible post-translational modification catalyzed by the sequential action of ubiquitin (E1), ubiquitin-conjugating (E2), and ubiquitin-ligating (E3) enzymes^6^, and reversed by deubiquitinating enzymes (DUBs)^7^. During viral infection, ubiquitin signaling fine-tunes cellular pathways to promote pathogen clearance while minimizing host damage^8^. However, viruses, such as IAV, which encodes a limited number of proteins and lacks its own ubiquitinating enzymes, have evolved to hijack the host ubiquitin system to expand the functional repertoire of viral proteins^9–11^. Consequently, the progression of IAV infection depends on a precarious balance between host-mediated degradation and viral-mediated subversion of the ubiquitin landscape.

Tripartite motif (TRIM) proteins are a large family of E3 ubiquitin ligases with key roles in innate immunity during viral infection^12–14^. Among them, TRIM21, which can assemble multiple ubiquitin chain types and undergoes dynamic auto-ubiquitination^15^, is unique as it bridges adaptive and innate immunity by functioning as a cytosolic antibody receptor^16^. This dual functionality positions TRIM21 as a key sensor and effector of intracellular immunity. However, the functional impact of TRIM21 remains paradoxical. While some studies suggest TRIM21 restricts replication though ubiquitin-mediated degradation of viral proteins and enhancement of innate signaling^17–22^, others indicate it may dampen antiviral responses to promote replication^23–28^. These observations indicate that TRIM21 does not operate as a simple on-off antiviral switch, but rather as a context-sensitive regulator whose activity may be shaped by expression level, infection stage, or interaction with parallel immune pathways. Moreover, our previous study showed that several TRIMs, including TRIM21, are differentially ubiquitinated during viral infection^29^, suggesting that it may function as a signal integrator rather than a unidirectional effector.

In this study, we sought to address the functional ambiguity surrounding TRIM21 during IAV infection. We demonstrate that TRIM21 exerts both proviral and antiviral effects through expression-dependent ubiquitin regulation of influenza nucleoprotein (NP) and innate signaling pathways. Furthermore, we identify PRKDC (DNA-PKcs) as a compensatory antiviral effector that is selectively engaged in the absence of TRIM21, revealing a hierarchical organization of host defenses. In summary, these findings define a context-dependent framework in which TRIM21 acts as a molecular rheostat that tunes IAV replication and innate immune output and reveals the TRIM21-PRKDC axis as a previously unappreciated layer of plasticity in host antiviral defense.

## RESULTS

### TRIM21 deficiency suppresses IAV replication

Since TRIM21 is an interferon-stimulated gene (ISG)^30^, we hypothesized that TRIM21-dependent innate immune signaling is activated during IAV infection to restrict viral replication. To test this hypothesis, we measured TRIM21 expression following infection with A/California/04/2009 (H1N1pdm). TRIM21 mRNA levels increased at 16 hours post infection (hpi) but declined at later time points (**Figure 1A**). In contrast, TRIM21 protein levels decreased during early infection and subsequently increased at later stages (**Figure 1B**). Moreover, both TRIM21 mRNA and protein levels increased in a dose-dependent manner with increasing multiplicity of infection (MOI) (**Figure 1A and S1A**). Our results show that TRIM21 expression is dynamically regulated in a time- and dose-dependent manner during infection.

**Figure 1.**
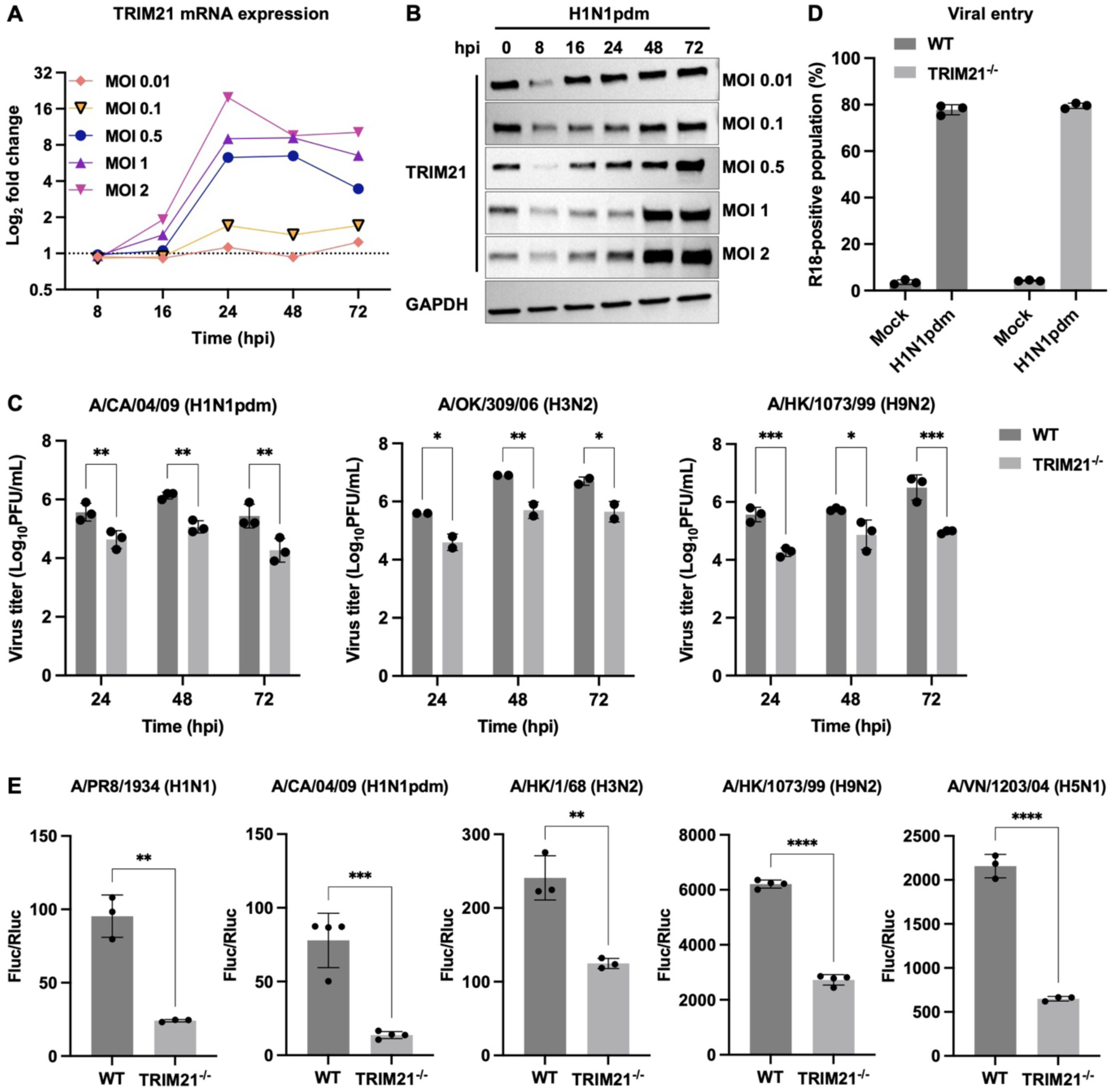
TRIM21 is required for efficient IAV replication. **(A and B)** A549 cells were infected with H1N1pdm at varying MOI (0.01, 0.1, 0.5, 1 and 2) and harvested at the indicated timepoints. **(A)** Total cellular RNA was isolated, and TRIM21 mRNA was analyzed by real-time PCR. The dotted line indicates fold change of 1. **(B)** Lysates prepared from infected cells were immunoblotted with antibodies against TRIM21. GAPDH was used as a loading control. **(C)** Viral titer in wild-type (WT) and TRIM21 knockout (TRIM21^−/−^) A549 cells infected with H1N1pdm, H3N2, or H9N2 (MOI = 0.01) at the indicated time points were measured by plaque assay. Data are shown as means of n = 2 or 3 ± standard deviations (SD). Two-way ANOVA was used to analyze data (*p < 0.05, **p < 0.01, ***p < 0.001). **(D)** Viral entry was assessed by flow cytometric analysis of R18-positive populations for 1 hpi. Data are shown as means of n = 3 ± standard deviations (SD). **(E)** Influenza polymerase reconstitution assay was performed in WT and TRIM21^−/−^HEK293T cells using RNP components from H1N1, H1N1pdm, H3N2, H9N2, or H5N1. Polymerase activity was normalized to Renilla luciferase. Data are shown as means of n = 3 or 4 ± standard deviations (SD). Two-tailed Student’s unpaired t-test was used to analyze data (**p < 0.01, ***p < 0.001, ****p < 0.0001).

To further investigate the function of TRIM21 during influenza virus infection, we first depleted TRIM21 using siRNA-mediated knockdown (siTRIM21) and infected the cells with IAV subtypes H1N1, H3N2, and H9N2. Viral production in siTRIM21 cells was reduced by 0.8 to 4.3 log units compared with non-targeting (NT) controls across all tested IAV strains (**Figure S1B and C**). Consistently, viral production was significantly reduced in two independent clones of TRIM21 knockout (TRIM21^−/−^) A549 cells compared with wild-type (WT) cells for all IAV strains tested (**Figure 1C and S1D**). These findings indicate that TRIM21 is involved in the modulation of influenza virus infection.

To determine the specific stage in the viral life cycle regulated by TRIM21, we first examined viral entry using previously described assays^31,32^. Infection of WT and TRIM21^−/−^ A549 cells with R18-labelled H1N1pdm showed no differences in viral entry and fusion (**Figure 1D and S1F**). By contrast, influenza polymerase activity was significantly reduced in TRIM21^−/−^ HEK293 cells for both low- and highly pathogenic influenza strains (**Figure 1E and S1E**). Together, these results demonstrate that TRIM21 does not affect viral entry but instead functions as a host factor required for efficient influenza virus replication.

### TRIM21 interacts with influenza NP through SPRY domain *in vivo* and *in vitro*

Given that TRIM21 modulates influenza polymerase activity, we next examined whether TRIM21 associates with the viral polymerase complex of A/Puerto Rico/8/1934 (PR8) using co-immunoprecipitation assay. TRIM21 was found to interact specifically with the NP (**Figure 2A**). In addition to PR8 NP, TRIM21 also interacted with NP from H1N1pdm and H9N2, consistent with the high degree of conservation of NP across diverse IAV strains^33^ (**Figure 2B**). Additionally, in uninfected cells, TRIM21 was predominantly localized in the cytoplasm, whereas H1N1pdm infection induced redistribution of TRIM21 to both the nucleus and cytoplasm (**Figure 2C**).

**Figure 2.**
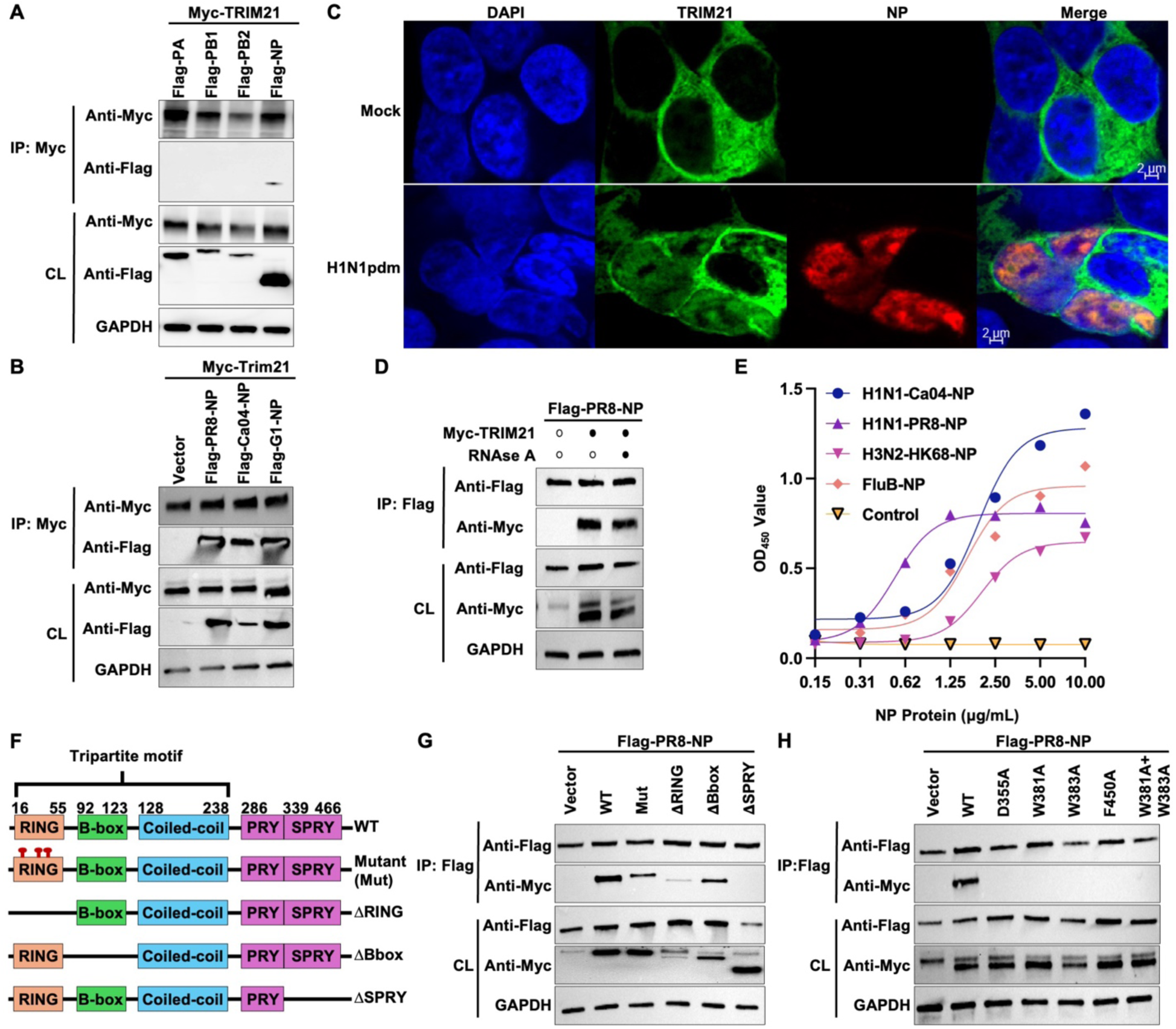
TRIM21 interacts with IAV and IBV NP. **(A)** HEK293T cells were transiently co-transfected with Myc-TRIM21 and Flag-tagged PA, PB1, PB2 or NP from A/Puerto Rico/8/1934 (PR8). Lysates were immunoprecipitated with anti-c-Myc magnetic beads, and immunoprecipitates (IP) were analyzed by immunoblotting with anti-Myc and anti-Flag antibodies. Transfected TRIM21 and viral proteins in cell lysates (CL) were confirmed by immunoblotting. **(B)** HEK293T cells were transiently co-transfected with Myc-TRIM21 and Flag-tagged NP from PR8, A/California/04/2009 (Ca04) or A/Quail/ Hong Kong/G1/97 (G1). Lysates were immunoprecipitated with anti-c-Myc magnetic beads and analyzed by immunoblotting. **(C)** Immunofluorescence imaging of TRIM21 and NP was performed in mock and H1N1pdm-infected A549 cells (MOI = 10). **(D)** HEK293T cells were co-transfected with Myc-TRIM21 and Flag-PR8-NP. Lysates were treated with or without RNAse A (50 mg/mL) for 10 minutes at 37°C prior to immunoprecipitation using anti-Flag M2 affinity beads. IP and CL were then analyzed by immunoblotting. **(E)** The binding affinities of H1N1-Ca04-NP (blue), H1N1-PR8-NP (purple), H3N2-HK68-NP (pink) and influenza B NP (orange) to TRIM21 were measured by ELISA. PBS was included as a negative control. **(F)** The schematic diagram of TRIM21 and its truncated mutants. TRIM21 contains a tripartite motif bearing an N-terminal RING domain, a B-box and coiled-coil domain. TRIM21 bears an PRYSPRY domain responsible for substrate interaction. Triad catalytic inactive mutant (C16A/C31A/H33W) is marked with red stalk. **(G and H)** HEK293T cells were transiently co-transfected with Flag-PR8-TRIM21 and Myc-TRIM21 constructs, including **(G)** WT, catalytic inactive mutant (Mut), or deletion mutants lacking the RING (ýRING), B-box (ýBbox) or SPRY (ýSPRY) domain, or **(H)** WT or point mutants within the SPRY domain. Lysates were immunoprecipitated with anti-Flag M2 affinity beads and then analyzed by immunoblotting.

Since NP is an RNA-binding protein^33^, we next assessed whether the TRIM21-NP interaction was RNA dependent. Treatment of co-immunoprecipitated lysates with RNase A did not disrupt the TRIM21-NP interaction (**Figure 2D**), indicating that the association is mediated by direct protein-protein interactions rather than bridged by viral RNA. To further validate this conclusion, we assessed TRIM21-NP *in vitro* binding using an ELISA-based assay. TRIM21 interacted with NP derived from both IAV and influenza B virus (IBV) (**Figure 2E**), confirming a conserved, RNA-independent interaction.

We next aimed to define the molecular nature of TRIM21-NP interaction. Flag-tagged-PR8-NP was co-expressed with either WT Myc-TRIM21, a catalytically inactive mutant harboring triple substitutions C16A/C31A/H33W within the RING domain (Mut), TRIM21 constructs lacking the RING (ΔRING), B-box (ΔBbox), or SPRY (ΔSPRY) deletion domains (**Figure 2F**). The catalytically inactive mutant results in complete loss of its catalytic activity^34^. Co-immunoprecipitation analyses revealed that deletion of the SPRY domain abolished NP binding, whereas catalytically inactive mutant and deletion of the RING or B-box domains did not (**Figure 2G**). Residues D355, W381, W383, and F450 within the SPRY domain of TRIM21 are known to mediate IgG Fc binding^16^. To test the importance of these residues for the interaction with NP, we mutated them to alanine. All of these substitutions abrogated NP binding (**Figure 2H**). These results show that the NP interacts with the SPRY domain of TRIM21.

### Dual ubiquitin linkage activity of TRIM21 shapes IAV replication

Previous studies have shown that ubiquitination of the NP is critical for efficient IAV replication^35–39^. We therefore investigated how TRIM21-mediated NP ubiquitination is regulated. To this end, we stably reconstituted WT TRIM21 (TRIM21^WT^), a catalytic inactive mutant (TRIM21^Mut^) or a SPRY domain-deficient variant (TRIM21^ΔSPRY^) in the TRIM21^−/−^ A549 and HEK293T cells (**Figure S2A**). Notably, TRIM21^Mut^ exhibited an aggregated localization pattern, consistent with its impaired catalytic function (**Figure S2B**). Following infection H1N1pdm virus, the expression of TRIM21^WT^, but not TRIM21^Mut^ and TRIM21^ΔSPRY^, restored the viral production in TRIM21^−/−^ A549 cells to a level comparable to that observed in WT cells (**Figure 3A**). A similar observation was also made for influenza polymerase activity (**Figure 3B**). These data demonstrate that both the E3 ubiquitin ligase activity and the SPRY domain of TRIM21 are necessary for modulating IAV replication.

**Figure 3.**
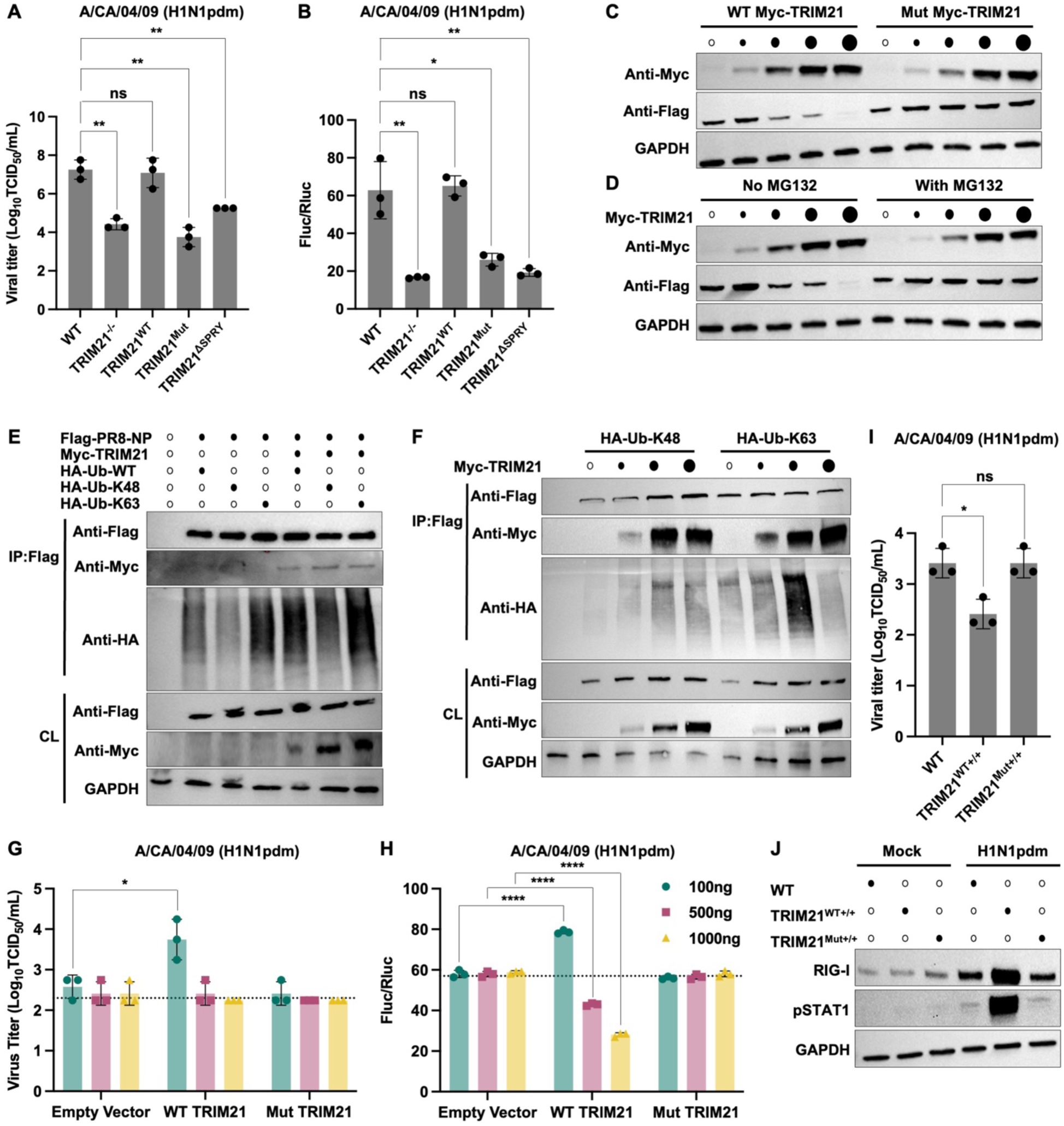
TRIM21 modulates IAV replication in an expression-dependent manner. **(A)** Viral titer in WT, TRIM21^−/−^, WT TRIM21-reconstitued (TRIM21^WT^), catalytic inactive mutant of TRIM21-reconstituted (TRIM21^Mut^) and SPRY domain-deficient TRIM21-reconstituted (TRIM21^ýSPRY^) A549 cells infected with H1N1pdm infection (MOI = 0.01) at 24 hpi were measured by TCID_50/_mL. Data are shown as means of n = 3 ± standard deviations (SD). Two-tailed Student’s unpaired t-test was used to analyze data (ns: not significant, **p < 0.01). **(B)** H1N1pdm influenza polymerase reconstitution assay in the indicated HEK293T cell lines. Polymerase activity was normalized to Renilla luciferase. Data are shown as means of n = 3 ± standard deviations (SD). Two-tailed Student’s unpaired t-test was used to analyze data (ns: not significant, *p < 0.05, **p < 0.01). **(C)** HEK293T cells were transiently co-transfected with Flag-PR8-NP and increasing amount of WT or Mut Myc-TRIM21. **(D)** HEK293T cells were transiently co-transfected with Flag-PR8-NP and Myc-TRIM21 and treated with or without 25 μM proteasomal inhibitor MG132 for 6 hours prior to harvest. **(C and D)** NP and TRIM21 expression were measured by immunoblotting using anti-Flag and anti-Myc antibodies, respectively. Increasing TRIM21 expression is indicated by progressively larger black dots; white dots indicate no TRIM21 transfection. **(E and F)** HEK293T cells were transiently co-transfected with Flag-PR8-NP, Myc-TRIM21, and HA-ubiquitin (WT versus linkage-specific constructs as indicated). In **(F)**, Myc-TRIM21 was transfected at increasing amount, denoted by progressively larger black dots. Cells were treated with MG132, followed by immunoprecipitation using anti-Flag M2 affinity beads. IP and CL were then analyzed by immunoblotting. Black and white dots indicate the presence or absence of plasmid transfection, respectively. **(G)** A549 cells were transiently transfected with empty vector control, WT Myc-TRIM21, or Mut Myc-TRIM21 at increasing amount and subsequently infected with H1N1pdm virus (MOI = 0.01). Viral titers at 24 hpi were determined by TCID_50/_mL. The dotted line indicates the mean viral titer of the empty vector control. Data are shown as means of n = 3 ± standard deviations (SD). Two-tailed Student’s unpaired t-test was used to analyze data (*p < 0.05). **(H)** H1N1pdm influenza polymerase reconstitution assay in HEK293T cells transiently transfected with empty vector control, WT Myc-TRIM21 or Mut Myc-TRIM21 at increasing amount, together with H1N1pdm RNP components and reporter constructs. Polymerase activity was normalized to Renilla luciferase. The dotted line indicates the mean polymerase activity of the empty vector control. Data are shown as means of n = 3 ± standard deviations (SD). Two-tailed Student’s unpaired t-test was used to analyze data (****p < 0.0001). **(I)** Viral titer in WT A549 cells, WT cells stably overexpressing TRIM21 (TRIM21^WT+/+^), or the catalytically inactive mutant (TRIM21^Mut+/+^) infected with H1N1pdm infection (MOI = 0.01) at 24 hpi were measured by TCID_50/_mL. Data are shown as means of n = 3 ± standard deviations (SD). Two-tailed Student’s unpaired t-test was used to analyze data (ns: not significant, *p < 0.05). **(J)** WT, TRIM21^WT+/+^, TRIM21^Mut+/+^ A549 cells were infected with H1N1pdm (MOI = 0.01). Total cell lysates were collected at 24 hpi and immunoblotted for RIG-I and phosphorylated STAT1 (pSTAT1). Cell lines are indicated by black dots.

We then aimed to determine whether TRIM21 regulates the expression and ubiquitination of NP via co-expression transfection assay. Increasing expression levels of WT TRIM21 but not the catalytic inactive mutant resulted in a dose-dependent decrease in NP level (**Figure 3C**). This effect was proteasome-dependent, as TRIM21-mediated NP degradation was abolished in the presence of the proteasome inhibitor MG132 (**Figure 3D**). These findings suggest that TRIM21 may function as either a pro- or antiviral factor depending on its expression level. K48-linked ubiquitination is classically associated with proteasomal degradation, whereas K63-linked ubiquitination has been reported to facilitate efficient IAV replication^9,10^. Given that TRIM21 is capable of catalyzing both K48- and K63-linked ubiquitination, we sought to elucidate the molecular basis underlying this dual pro- and antiviral function by *in vitro* ubiquitination assay. Our results showed that TRIM21 catalyzed NP ubiquitination through both K48- and K63-linked polyubiquitin chains (**Figure 3E**). Notably, NP ubiquitin linkage specificity was TRIM21 expression-dependent, with intermediate TRIM21 levels favoring K63-linked polyubiquitination, whereas higher expression shifted ubiquitination toward K48 linkage (**Figure 3F**). Functionally, this linkage switch correlated with viral outcomes. WT-level TRIM21 expression restored viral replication to baseline (**Figure 3A-B**), and moderate TRIM21 expression enhanced viral replication (**Figure 3G-H)**. In contrast, high TRIM21 expression suppressed viral replication in a catalytic activity-dependent manner (**Figure 3G-H)**, coinciding with increased K48-linked ubiquitination and NP degradation. Together, these results suggest that TRIM21 differentially regulates NP fate and viral replication through linkage-specific ubiquitination.

As an ISG, TRIM21 expression was induced by multiple innate immune signaling stimuli (**Figure S2C**) and increased in a dose-dependent manner following interferon (IFN) treatment (**Figure S2D**). Consistently, overexpression of WT TRIM21 but not the Mut Trim21 augmented interferon-stimulated response element (ISRE) activity (**Figure S2E**), indicating a positive feedback loop with innate immune signaling. We therefore hypothesized that, in addition to the TRIM21-mediated proteasome-dependent NP degradation (**Figure 3D**), enhanced innate immune responses contribute to the antiviral effect observed at high TRIM21 expression levels. Because transient transfection efficiency in A549 cells is limited, we generated A549 cells stably overexpressing either TRIM21 (TRIM21^WT+/+^) or the catalytically inactive mutant TRIM21 (TRIM21^Mut+/+^) (**Figure S2F**) and infected them with H1N1pdm virus. TRIM21^WT+/+^ resulted in reduced viral production compared with WT control cells, whereas no significant effect was observed in TRIM21^Mut+/+^ (**Figure 3I**). This reduction was not attributable to impaired viral entry (**Figure S2G-H**), but due to enhanced innate immune response observed in WT TRIM21^WT+/+^ during infection (**Figure 3J**). These results suggest that high TRIM21 expression suppresses IAV replication by simultaneously mediating proteasomal-dependent NP degradation and amplifying host antiviral signaling.

### TRIM21-PRKDC ubiquitination axis tunes antiviral signaling

To define how TRIM21 deficiency reshapes host responses during influenza virus infection, we integrated transcriptomic and quantitative proteomic analyses of WT and TRIM21^−/−^ A549 cells under mock and H1N1pdm infection conditions. Unexpectedly, differential expression analysis revealed a pronounced enrichment of ISGs (*IFI44L*, *IFI6*, *MX1*, *HERC6, and EIF2AK2*) in infected TRIM21^−/−^ cells compared with WT cells (**Figure S3A**). Additionally, Gene Ontology (GO) analysis of genes upregulated in infected TRIM21^−/−^ cells showed a significant antiviral and innate immune pathways activation (**Figure S3B**). Notably, uninfected TRIM21^−/−^ cells exhibited an intrinsically lower basal antiviral transcriptional profile than the WT cells (**Figure S3C and Table S1**). Consistently, proteomics analysis showed that although basal ISG expression was diminished in TRIM21^−/−^ cells, infection induced ISG protein expression to levels comparable to those observed in WT cells (cluster 2 in **Figure 4A**), despite reduced virus production in the absence of TRIM21 (**Figure 1C**). Similarly, immunoblot analysis validated that both viral infection and polyinosinic-polycytidylic acid (PolyIC) stimulation elicited WT-comparable innate immune responses in TRIM21^−/−^ cells (**Figure 4B and S3D**). Collectively, these results indicate that TRIM21 deficiency lowers the threshold for innate immune activation, resulting in a disproportionately robust IFN-stimulated response upon immune challenge.

**Figure 4.**
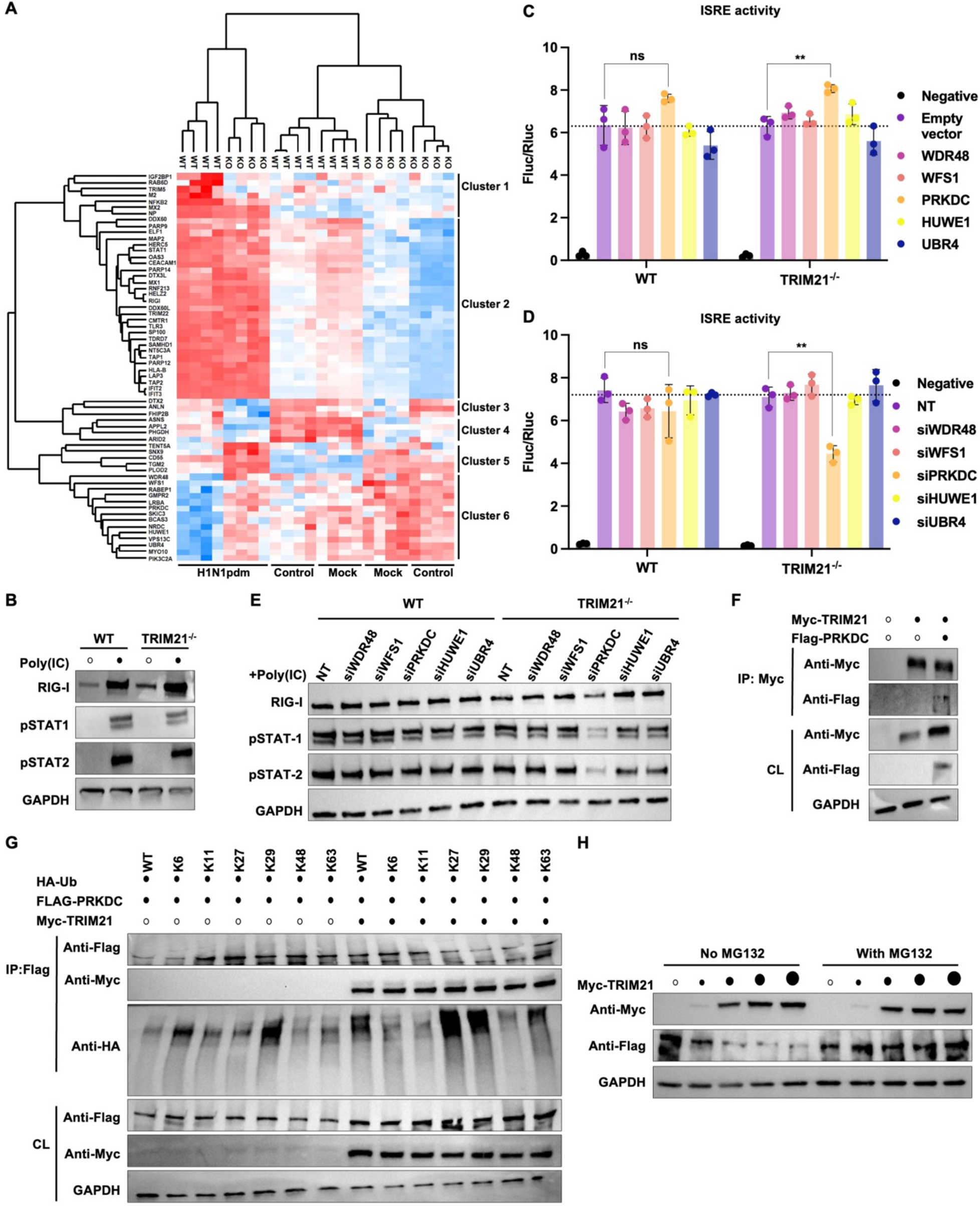
TRIM21 negatively regulates PRKDC to tune innate immune output. **(A)** Heatmap showing the relative abundance of proteins identified as statistically significant (p < 0.05) by two-way ANOVA across genotypes (WT or TRIM21^−/−^), conditions (basal control, mock- or H1N1pdm-infected), and their interaction. Protein abundance was calculated using maximum label-free quantification per protein per sample, normalized with z-score. Only proteins significant in all three comparisons (genotype, infection and interaction) are shown. Colors indicate upregulated (red) or downregulated (blue) proteins. Experiment was performed in four replicates for each condition. The cells were harvested at 48 hpi (MOI 0.01), the time point at which viral production differed most markedly between the cells **(B)** WT and TRIM21^−/−^ A549 cells were treated with polyIC for 24 hours, and cell lysates were immunoblotted for RIG-I, pSTAT1, and phosphorylated STAT2 (pSTAT2). **(C and D)** HEK293T cells were transiently transfected with the indicated **(C)** plasmids or **(D)** siRNAs, together with an ISRE-driven firefly luciferase reporter and a Renilla luciferase control. At 24 hours post-transfection, cells were treated with IFN-α for an additional 24 hours prior to measurement of luciferase activities. Data are shown as means of n = 3 ± standard deviations (SD). Two-tailed Student’s unpaired t-test was used to analyze data (ns: not significant, **p < 0.01). The dotted line indicates the mean ISRE activity in empty vector or non-targeting (NT) control. **(E)** The indicated proteins were depleted by siRNA in both WT and TRIM21^−/−^ A549 cells, followed by polyIC treatment. Lysates were immunoblotted for RIG-I, pSTAT1, and pSTAT2. **(F)** HEK293T cells were transiently co-transfected with Myc-TRIM21 and Flag-PRKDC. Lysates were immunoprecipitated with anti-c-Myc magnetic beads and analyzed by immunoblotting. Black and white dots indicate the presence or absence of plasmid transfection, respectively. **(G)** HEK293T cells were transiently co-transfected with Flag-PRKDC, Myc-TRIM21, and HA-ubiquitin (WT versus linkage specific constructs as indicated). Cells were treated with MG132, followed by immunoprecipitation using anti-Flag M2 affinity beads. IP and CL were then analyzed by immunoblotting. Black and white dots indicate the presence or absence of plasmid transfection, respectively. **(H)** HEK293T cells were transiently co-transfected with Flag-PRKDC and increasing amount of WT Myc-TRIM21 and treated with or without proteasomal inhibitor MG132. PRKDC and TRIM21 expression were measured by immunoblotting using anti-Flag and anti-Myc antibodies, respectively.

To elucidate the mechanism enabling TRIM21^−/−^ cells to achieve WT-level ISG induction despite their lower basal state, we examined a subset of proteins that displayed sustained or elevated expression in TRIM21^−/−^ cells during infection (cluster 6 in **Figure 4A**). This subset of proteins are functionally important for ubiquitin signaling, innate immunity, and stress responses, such as WDR48^40,41^, WFS1^42^, PRKDC^43,44^, HUWE1^45–47^, and UBR4^45,48,49^. Among these factors, transient PRKDC overexpression robustly enhanced ISRE activity in TRIM21^−/−^ cells (**Figure 4C**). Conversely, siRNA-mediated silencing of these candidates revealed that among the tested genes only depletion of PRKDC significantly attenuated ISRE activity in TRIM21^−/−^ cells (**Figure 4D and S3E**). This dependency was further validated in A549 cells, where PRKDC silencing significantly reduced polyIC-induced innate immune responses specifically in TRIM21^−/−^ cells (**Figure 4E and S3F**). Of note, PRKDC is a serine/threonine protein kinase that acts as a molecular sensor for DNA damage, and has been implicated in regulating innate immune response during DNA virus infection^43,50,51^. Together, these data identify PRKDC as a key compensatory regulator that sustains innate immune signaling in the absence of TRIM21, revealing a previously unrecognized TRIM21-PRKDC axis that buffers antiviral immunity during influenza infection.

Given that TRIM21 functions as an E3 ubiquitin ligase, we aimed to determine whether TRIM21 regulated the expression and ubiquitination of PRKDC. We first confirmed that PRKDC undergoes ubiquitination (**Figure S3G**). Furthermore, co-immunoprecipitation assay demonstrated an interaction between TRIM21 and PRKDC (**Figure 4F**). Linkage-specific ubiquitination analyses revealed that TRIM21 catalyzed K27- and K29-linked polyubiquitination of PRKDC (**Figure 4G**), which in turn promoted proteasome-dependent degradation of PRKDC (**Figure 4H**). These data indicate that TRIM21 acts as a negative regulator of PRKDC. The absence of TRIM21 would increase the protein stability of PRKDC, thereby lowering the activation threshold for antiviral gene induction during influenza infection.

### PRKDC functions as an antiviral restriction factor in the absence of TRIM21

To determine the functional significance of PRKDC-TRIM21 axis in regulating influenza virus replication, we assessed the effect of PRKDC depletion on viral production in WT and TRIM21^−/−^cells. Knockdown of PRKDC by siRNA did not affect viral titers in WT cells. In contrast, PRKDC knockdown in TRIM21^−/−^ cells led to significant increase in viral titers, indicating that PRKDC becomes a critical antiviral factor specifically when TRIM21 is absent (**Figure 5A**). The increase of viral production was not attributable to enhanced viral entry (**Figure S4A-B**) but associated with elevated viral polymerase activity (**Figure 5B**). To further assess whether PRKDC influences vRNP nuclear export, we performed RNA fluorescence in situ hybridization (FISH) using probes that were specific to PB2 and NP vRNA. PRKDC depletion did not alter the nuclear export of vRNPs. At the same time, we found that PB2 and NP vRNA abundance was markedly reduced in TRIM21^−/−^ cells compared with WT cells (**Figure 5C)**, consistent with the reduction of viral titer and polymerase activity (**Figure 1C and E**). PRKDC knockdown restored the PB2 and NP vRNA levels in TRIM21^−/−^ cells to near WT levels **(Figure 5C)**, mirroring the observed increase in viral polymerase activity (**Figure 5B**). These data indicate that PRKDC restricts influenza virus replication by suppressing viral polymerase activity in TRIM21-deficient cells, revealing a context-dependent antiviral function that is masked in the presence of TRIM21.

**Figure 5.**
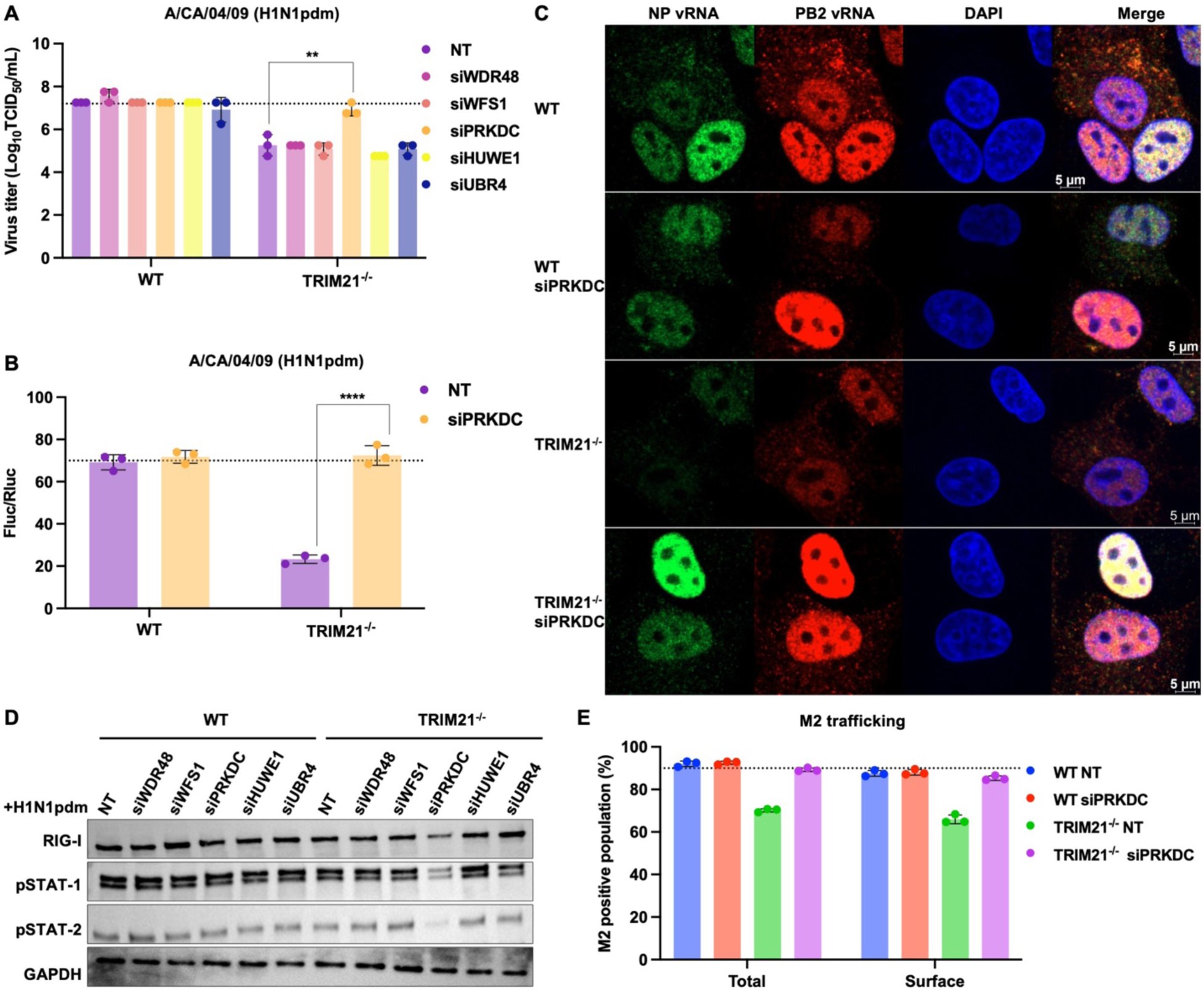
PRKDC is essential for the compensatory antiviral response in TRIM21-deficient cells. **(A)** WT and TRIM21^−/−^ A549 cells were transfected with non-targeting siRNA (NT) or siRNA targeting the indicated protein of interest, followed by infection with H1N1pdm (MOI = 0.01). Viral titers at 24 hpi were determined by TCID_50/_mL. The dotted line indicates the mean viral titer of NT-treated WT A549 cells. Data are shown as means of n = 3 ± standard deviations (SD). Two-tailed Student’s unpaired t-test was used to analyze data (**p < 0.01). **(B)** WT and TRIM21^−/−^ HEK293T cells were transfected with NT or PRKDC-targeting siRNA and subjected to an H1N1pdm influenza polymerase reconstitution assay. Polymerase activity was normalized to Renilla luciferase. The dotted line indicates the mean polymerase activity in NT-treated WT A549 cells. Data are shown as means of n = 3 ± standard deviations (SD). Two-tailed Student’s unpaired t-test was used to analyze data (****p < 0.0001). **(C)** RNA-FISH analysis in indicated PR8-infected A549 cells (MOI = 10). **(D)** The indicated proteins were depleted by siRNA in WT and TRIM21^−/−^A549 cells, followed by H1N1pdm infection (MOI = 0.01). Lysates were immunoblotted for RIG-I, pSTAT1, and pSTAT2. **(E)** WT and TRIM21^−/−^ A549 cells were transfected with NT or PRKDC-targeting siRNA and subsequently infected with H1N1pdm (MOI = 10). Total and surface expression of viral M2 protein were quantified by flow cytometry. The dotted line indicates the mean M2 expression in NT-treated WT A549 cells. Data are shown as means of n = 3 ± SD.

As we have identified PRKDC as a key compensatory regulator sustaining innate immune signaling in the absence of TRIM21, we next assessed whether this dependency was recapitulated during viral infection. PRKDC depletion in H1N1pdm-infected TRIM21^−/−^ cells resulted in reduced expression of RIG-I and diminished phosphorylation of STAT-1 and STAT-2 (**Figure 5D**). This result suggests that TRIM21^−/−^ cells develop a functional dependency on PRKDC to sustain antiviral signaling. In other words, our data reveals that PRKDC is an essential driver of the compensatory innate immune response in the absence of TRIM21.

Given that both TRIM21 and PRKDC have been implicated in autophagy^52–56^, and that viral proteins can be degraded by autophagic pathways^31,32,57^, we then evaluated whether autophagy contributed to the phenotypes observed. Beclin-1, p62 and LC3-II accumulation were comparable across all conditions, suggesting that neither PRKDC nor TRIM21 depletion led to altered autophagic flux (**Figure S4C-D**). Consistent with this, confocal microscopy revealed no significant differences in colocalization of viral M2 with the lysosomal marker LAMP1 across all conditions (**Figure S4F-G**). Moreover, viral trafficking remained unchanged across the conditions, and viral protein expression (M2 and HA) in PRKDC-depleted TRIM21^−/−^ was restored to levels comparable to those in WT (**Figure 5E and S4E**). Notably, no physical interaction between PRKDC and M2 was observed in either WT or TRIM21^−/−^ cells, suggesting that PRKDC does not directly modulate M2 protein expression (**Figure S4H**). Collectively, these data indicate that TRIM21 deficiency does not inherently drive autophagic degradation of the virus and that the restoration of viral protein expression observed in PRKDC-depleted TRIM21^−/−^ cells is not attributable to altered autophagy. Therefore, the recovery of viral titer in PRKDC-depleted TRIM21^−/−^ cells is driven by the overall surge in polymerase activity and replication, rather than a primary change in protein stability.

### TRIM21 orthologs preserve NP binding and modulation of antiviral signaling

Recurrent positive selection is a hallmark of host-pathogen arms races^58^. Nevertheless, site-model analysis revealed no statistically significant evidence of positive selection on TRIM21 in four distinct mammalian clades (**Figure S5A**), indicating that its evolution has been largely constrained by purifying selection. Furthermore, sequence analysis showed that TRIM21 is highly conserved among primates, with modestly increased sequence divergence observed in non-primate species, including rodents, carnivores, and ungulates (**Figure 6A**). Multiple sequence alignments further demonstrated high sequence conservation of residues critical for E3 ligase activity within the RING domain^34^, as well as residues within SPRY domain previously implicated in antibody interactions^16^ (**Figure S5B-C**). Consistent with this, co-immunoprecipitation assays demonstrated that all tested TRIM21 orthologs retained the ability to interact with influenza NP (**Figure 6B**). Moreover, expression of each TRIM21 ortholog recapitulated the human TRIM21 phenotype, exhibiting dose-dependent modulation of both influenza polymerase activity (**Figure 6C**) and ISRE activity (**Figure 6D**). Together, these results indicate that the core molecular functions of TRIM21 are preserved across mammals.

**Figure 6.**
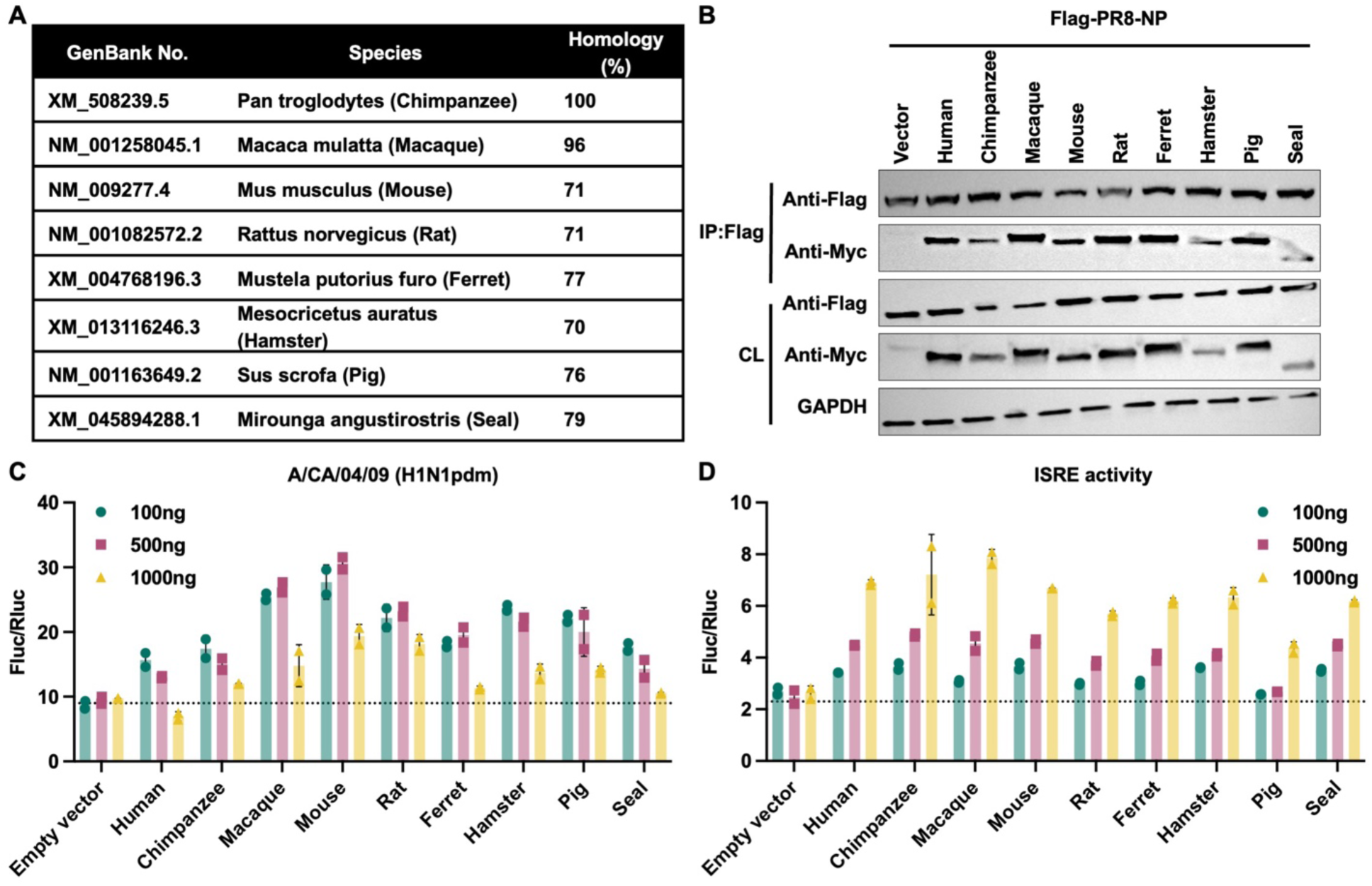
TRIM21 orthologs retain NP binding with dose-dependent functional output. **(A)** Sequence homology comparison between human TRIM21 and TRIM21 orthologs from representative species. **(B)** HEK293T cells were transiently co-transfected with Flag-PR8-NP and Myc-TRIM21 orthologs from the indicated species. Lysates were immunoprecipitated with anti-Flag M2 affinity beads and analyzed by immunoblotting. **(C)** H1N1pdm influenza polymerase reconstitution assay in HEK293T cells transiently transfected with empty vector control or Myc-TRIM21 orthologs at increasing amount, together with H1N1pdm RNP components and reporter constructs. Polymerase activity was normalized to Renilla luciferase. The dotted line indicates the mean polymerase activity in empty vector control. Data are shown as means of n = 2 ± standard deviations (SD). **(D)** HEK293T cells were transiently transfected with empty vector control or Myc-TRIM21 orthologs at increasing amount, together with an ISRE-driven firefly luciferase reporter and a Renilla luciferase control. At 24 hours post-transfection, cells were treated with IFN-α for an additional 24 hours prior to measurement of luciferase activities. The dotted line indicates the mean ISRE activity in empty vector control. Data are shown as means of n = 2 ± standard deviations (SD).

## DISCUSSION

The clinical and technological impact of TRIM21, from its identification as a diagnostic autoantigen^59^ to its revolutionary application in Trim-Away mediated protein depletion^60–62^, has positioned it as a central node of intracellular immunity. Despite this growing interest, the functional role of TRIM21 during viral infection remains incompletely understood, with seemingly conflicting findings describing it as either antiviral or proviral^63^. This paradox underscores a major gap in our understanding and suggests that the regulatory logic of TRIM21-mediated immunity is far more complex than currently appreciated.

A key conceptual advance of this study is the identification of TRIM21 as a context-dependent regulator of IAV infection rather than a uniformly antiviral or proviral factor. We demonstrate that TRIM21 functions as a molecular rheostat, in which its expression level and ubiquitin-dependent activity tune the balance between viral replication and innate immune output. This framework provides a mechanistic explanation for previously conflicting reports of TRIM21 function by suggesting that divergent phenotypes arise from differences in cellular context, infection stage, or TRIM21 abundance. By integrating direct effects on viral components with modulation of antiviral signaling pathways, TRIM21 operates as a regulatory hub rather than a simple effector. Furthermore, the requirement for its catalytic activity across both proviral and antiviral outcomes underscores ubiquitin signaling as a dynamic, information-encoding system capable of generating graded immune responses. Of note, TRIM family members exhibit highly heterogeneous, cell-type-specific expression patterns and extensive alternative splicing^64–66^, features that may similarly confer context-dependent regulatory functions. These observations raise the broader possibility that expression-tuned regulation, rather than binary restriction, may be a common and yet underappreciated organizing principle within the TRIM protein family.

Another highlight of this study is the identification of PRKDC functions as a compensatory restriction factor that becomes functionally relevant only when TRIM21-mediated regulation is lost. This selective requirement underscores functional redundancy and hierarchical organization within innate immune networks. Mechanistically, our findings also extend emerging evidence that PRKDC contributes to virus sensing and IFN regulation^43,44,67^. The mechanism by which TRIM21 navigates its broad repertoire of substrates during infection, including PRKDC, viral NP, and various autophagy and innate immune components^68^, remains a compelling frontier as it will profoundly impact viral infection. We propose that this selectivity may be governed by the differential binding affinities or by intrinsic turnover kinetics of the ligase itself. Unlike the rapid turnover of TRIM5α, the relative stability of TRIM21^69^ may enable it to function as a longer-term signal integrator that buffers immune activation and prevents premature engagement of secondary restriction pathways. Consequently, the proviral effects attributed to TRIM21 may reflect its role as a gatekeeper that limits activation of secondary, more potent restriction pathways such as PRKDC. This homeostatic restraint has been observed in the recruitment of USP18 by STAT2^70^ and the role of NLRC5^71^ in preventing excessive Type I IFN signaling. However, while these examples represent negative feedback loops, the TRIM21–PRKDC axis represents a paradigm shift. Unlike temporal fine-tuning or competitive ubiquitination of individual sensors seen with TRIM38^72^, TRIM25, and RNF125^73^, we demonstrate that E3 ligase “deficiency” can act as a programmed trigger. Here, the loss of a primary ligase activates a latent, secondary defense architecture that ensures immune continuity.

From an evolutionary perspective, TRIM family proteins such as TRIM5α have evolved under strong virus-driven selective pressures, with species lineage-specific expansions and recurrent positive selection shaping its antiviral activity^45,74^. In contrast, our evolutionary and functional analyses indicate that TRIM21 occupies a fundamentally different niche and experiences strong purifying selection across mammalian lineages, consistent with the preservation of core molecular functions rather than rapid diversification driven by virus-specific arms races. Concordantly, our data demonstrate that TRIM21 ubiquitin ligase activity is required for both its pro- and antiviral functions, underscoring a conserved, ubiquitin-dependent regulatory role that is distinct from TRIM5α, in which ubiquitin-proteasomal system is dispensable for antiviral activity^75^. Together, these findings position TRIM21 not as a classical restriction factor, but as a conserved immune rheostat that tunes antiviral responses across distinct mammalian species. Notably, whether this evolutionary constraint is maintained in non-mammalian or high-temperature influenza reservoirs such as bats^76^ remains an important open question with implications for host-specific regulation of influenza virus infection.

In summary, our study establishes TRIM21 as an expression-sensitive regulator of IAV replication that integrates ubiquitin signaling with innate immune control (**Figure 7**). By coupling NP ubiquitination to immune amplification while restraining PRKDC-dependent antiviral signaling, TRIM21 sets a tunable antiviral threshold that balances viral replication and host defense. Importantly, TRIM21 does not operate in isolation as its absence forces the cell to “re-wire” its proteome that engages secondary antiviral safeguards. This layered organization of host defense highlights the robustness of innate immune networks and underscores the need for caution when targeting E3 ligase pathways therapeutically, as perturbing primary regulators may unintentionally destabilize critical compensatory mechanisms.

**Figure 7.**
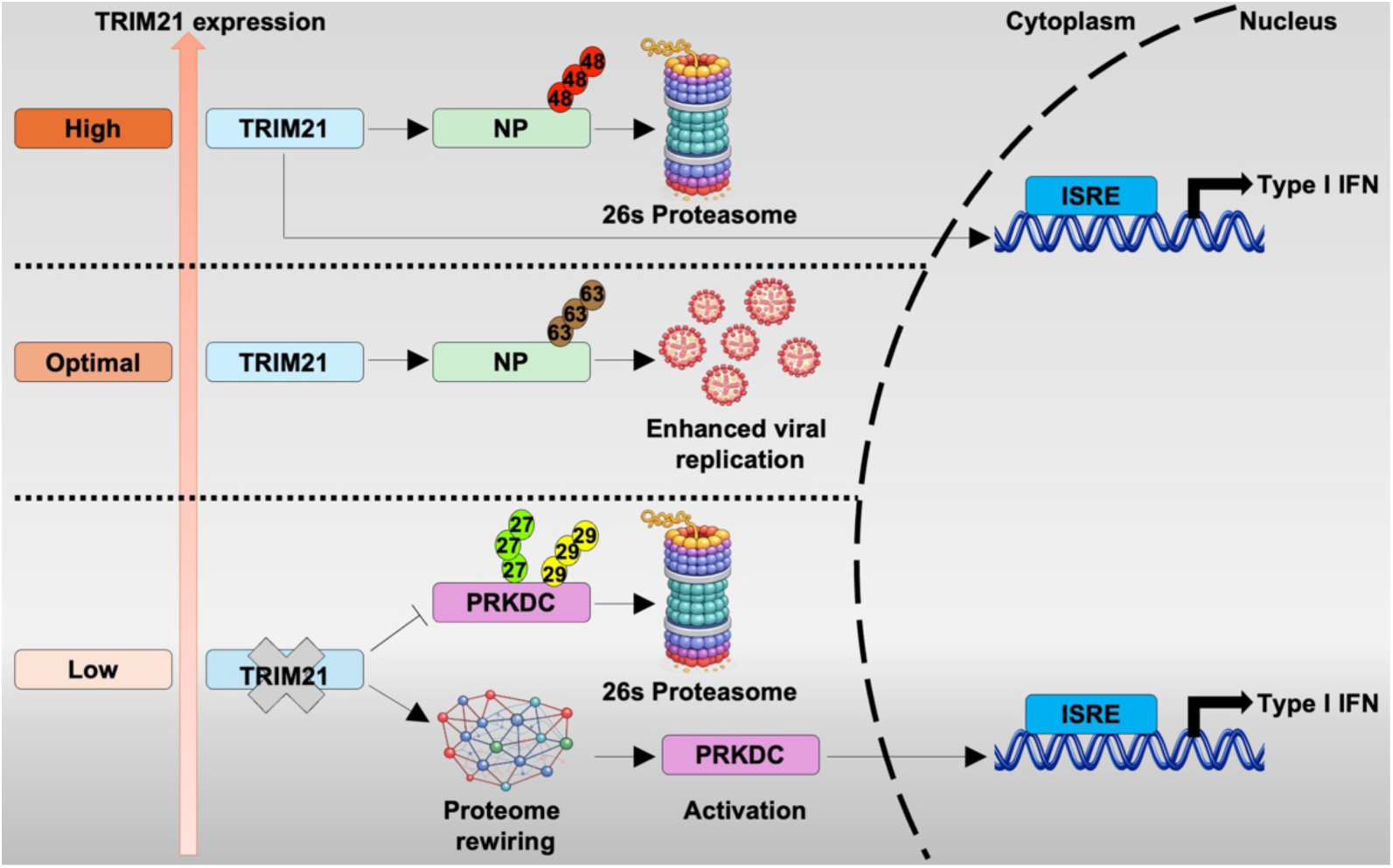
Schematic diagram of TRIM21 establishes a tunable antiviral threshold during IAV infection. Upon infection, TRIM21 interacts with NP and promotes K63-linked polyubiquitination, facilitate influenza polymerase activity and supporting viral replication under optimal expression levels. When the TRIM21 expression is elevated, it shifts toward promoting K48-linked polyubiquitination of NP, targeting NP for proteasomal degradation and amplifying innate immune signaling, thereby limiting viral replication. In the absence of TRIM21, systematic rewiring of cellular proteome is triggered and PRKDC expression is stabilized. This compensatory activation of PRKDC-driven antiviral pathway engages secondary antiviral safeguards to counteract viral infection.

## ACKNOWLEDGEMENTS

We thank the Roy J. Carver Biotechnology Center at the University of Illinois at Urbana-Champaign for assistance with RNA sequencing. We are grateful to the Institute of Genomic Biology Core Facility at the University of Illinois at Urbana-Champaign for providing access to the LSM880 microscope system. UBR4 plasmids were a gift from Prof. Yong Tae Kwon at Seoul National University. This work was supported by the Carl R. Woese Institute for Genomic Biology Postdoctoral Fellowship (H.L.), NIH R35GM137961 (L.R.), Wellcome Trust (220776/Z/20/Z and 223107/Z/21/Z to S.S), the Biotechnology and Biological Sciences Research Council (BB/Y000307/1 to S.S.), Vallee Scholars Program (N.C.W.), the Searle Scholars Program (N.C.W.), and Howard Hughes Medical Institute Emerging Pathogens Initiative (N.C.W.).

## AUTHOR CONTRIBUTIONS

Q.W.T., S.S., and N.C.W. conceived and designed the study. Q.W.T., H.L., P.P., X.C., V.R., K.J.M., J.J.H., J.R.-C., J.Z., Y.W.H., E.K.S., H.M.T., C.B.B., and L.R. assisted with experiments. Q.W.T., S.S., and N.C.W. wrote the paper and all authors reviewed and/or edited the paper.

## METHODS

### Cells lines and virus stocks

The following cell lines were used in this study: A549 (human; sex: male; ATCC-CCL-185), HEK293T (human; sex: unspecified) and MDCK (Canis familiaris; sex: unspecified; ATCC-CCL-34) were obtained from ATCC and gender of the cell line was not a consideration in the study. Cells were maintained in Dulbecco’s Modified Eagle Medium (DMEM; GIBCO) supplemented with 10% Fetal Bovine Serum (FBS; Thermo Fisher Scientific) and 1% Penicillin/Streptomycin (P/S; Thermo Fisher Scientific) at 37°C in a humidified incubator under 5% CO_2_ supply. The following virus strains were used in this study: A/Puerto Rico/8/1934 (H1N1), A/California/07/2009 (H1N1 pdm), A/Oklahoma/309/06 (H3N2) and A/Hong Kong/1073/99 (H9N2). Influenza viruses were propagated in Madin Darby Canine Kidney (MDCK) cells in DMEM supplemented with 1 μg/mL Tosyl Phenylalanyl Chloromethyl Ketone (TPCK)-treated trypsin (Thermo Scientific). Supernatants from the infected MDCK cells were harvested three days post-infection. In order to maintain the homogeneity of the viruses, they were propagated with limited passage number and seed stocks of viruses were prepared for future propagation. Virus stocks were aliquoted and stored at −80°C, and the viral titers were measured using plaque assay.

### Generation of cell lines

TRIM21 knockout cells were generated using single-guide RNA (sgRNA) sequences (TRIM21_A: 5-CACCGGAAACACCGTGACCACGCCA-3’ and 5’-AAACTGGCGTGGTCACGGTGTTTCC-3’. These were cloned into lentiCRISPR v2 (Addgene #52961) as previously described^77^. Cells were then infected by the lentivirus in the presence of polybrene (Sigma-Aldrich) for 72 hours. Puromycin selection was performed following the lentivirus transduction at a concentration of 1.4 μg/mL in A549 and 0.8 μg/mL in HEK293T cells for 72 hours. Single clones were picked by limiting dilution and confirmed by western blotting using specific antibodies. The stable cell lines expressing either WT or mutant TRIM21 were generated using retroviral vector pCHMES-IRES-Hygromycin, selected following a 2-week period in the presence of 1.6 mg/mL in A549 and 0.1 mg/mL in HEK293T cells. Single clones were picked by limiting dilution and confirmed by western blotting using specific antibodies. Selected positive clones were maintained thereafter in the same selection medium. All siRNAs used in this work were purchased from Dharmacon as SMARTpool ON-TARGET plus siRNA. Cells were seeded in 24-well plates with the seeding density of 0.02 x 10^6^ cells/well one day in advance. siRNAs mixed with DharmaFECT1 were added to cells in DMEM without antibiotics and incubated at 37°C for 48 hours. Gene knockdown efficiency was quantified by qPCR.

### Virus infection

Cells were seeded in 24-well plates with the seeding density of 0.1 x 10^6^ cells/well one day in advance. Cells were infected with viruses at a multiplicity of infection (MOI) of 0.01 for multi-cycle infection assay. After 1 hour of adsorption, the viral inoculum was removed, and the infected cells were washed with Phosphate-buffered Saline (PBS; Thermo Fisher Scientific) and maintained in DMEM supplemented with 0.2 μg/mL TPCK-treated trypsin at 37°C. Supernatants, cell lysates, and total RNA were harvested at indicated post infection time points, cleared by centrifugation and stored at −80°C until further analysis.

### Plaque assay

MDCK cells were seeded in 12-well plates with the seeding density of 0.5 x 10^6^ cells/well one day in advance. The following day confluent MDCK cells were washed with PBS and then incubated with 500 μl of the ten-fold serially diluted infectious media for 1 hour at 37°C. Afterwards, the viral inoculum was removed. Infected cells were washed twice with PBS and overlaid with DMEM containing 1% agarose and 1 μg/mL TPCK-treated trypsin. The plates were incubated in inverted position at 37°C for three days. Cells were fixed with 4% formaldehyde in PBS overnight. Plaques were visualized by staining the plates with 1% crystal violet in 20% ethanol for 15 minutes. Plaques were counted and recorded after drying the plates.

### Viral entry assay

Cells were infected with virus labelled with the fluorescent probe octadecyl rhodamine B Chloride (R18). R18 is self-quenching at high concentrations and shows no fluorescence signal when prelabelled on virus. However, as R18-labelled virus fuses with unlabelled cellular membranes, R18-dilution occurs leading to increased fluorescence intensity. This increased fluorescence signal was quantified as a measure of viral fusion using flow cytometry. Pre-labelling of virus with R18 was executed by incubation of virus or control media with 1:1000 2× R18 dye (Thermo Fisher) for 1 hour on ice. Residual dye was removed from the labelled virus by ultracentrifugation on a 25% sucrose gradient at 28000 rpm for 1 hour. After resuspension of the virus on ice, cells were incubated with the virus at MOI 10 (as determined prior to centrifugation) for 1 hour at 4°C followed by 1 hour at 37°C. Afterwards the cells were washed, trypsinised and fixed (4% PFA) for analysis on flow cytometry (Novocyte Opteon, Agilent). The %-R18+ population was analyzed using FlowJo^TM^ v10.8 Software (BD Life Sciences).

### Influenza virus polymerase activity assay

Dual luciferase activity reporter assays were performed to compare the polymerase activity of RNP complexes. RNP complex expression plasmids composed of PB2, PB1, PA (125 ng each) and NP (250 ng), together with pYH-Luci reporter plasmid (125 ng) and Renilla reporter plasmid (10 ng) were co-transfected into HEK293T cells. At 24 hours after transfection, cells were lysed, and the luciferase activity was measured using the Dual-Luciferase reporter assay system (Promega). Measurements were acquired by a BioTek synergy HTX multimode reader (Agilent).

### ISRE luciferase assay

For luciferase assays, HEK293T cells were seeded in a 24-well plate at a density of 100,000 cells per well. The next day, cells were transiently transfected with pRL-TK and ISRE-Luc reporter plasmids along with the indicated plasmids for 24 hours. Cells were then treated overnight with 2000 U/mL of universal Type I IFN (PBL Bioscience) and lysed at 24 hours post treatment using Passive Lysis Buffer (Promega). Samples were processed and luciferase activity was measured using the Dual-Luciferase Assay System (Promega) according to the manufacturer’s instructions. Measurements were acquired by BioTek synergy HTX multimode reader (Agilent). Firefly luciferase values were normalized to Renilla luciferase values.

### Enzyme-linked immunosorbent assay (ELISA)

Nunc Maxisorp ELISA plates (Thermo Fisher Scientific) were coated with 100 μL of TRIM21 protein at 1 μg/mL in a 1 x PBS solution overnight at 4°C. The plates were washed 3 times the next day with 1 x PBS supplemented with 0.05% Tween 20 and blocked with 100 μL of ChonBlock Blocking/Sample Dilution ELISA Buffer (Chondrex, Inc.) for 1 hour at room temperature. NP proteins or PBS were serially diluted 1:2 starting at 10 μg/mL and incubated for 2 hours at 37°C. The plates were then washed 3 times and incubated with pooled human plasma samples (positive for NP protein) diluted 1:50 for 2 hours at 37°C. The plates were further washed 3 times and incubated with horseradish peroxidase (HRP)-conjugated goat anti-human IgG antibody (GE Healthcare) diluted 1:5000 for 1 hour at 37°C. After 6 washes with 1 x PBS supplemented with 0.05% Tween 20, 100 μL of 1-Step TMB ELISA Substrate Solution (Thermo Fisher Scientific) was added to each well. After a 10 minutes incubation, the reaction was stopped with 50 μL of 2 M H_2_SO_4_ solution, and absorbance values were measured at 450 nm using a BioTek synergy HTX multimode reader (Agilent).

### Proteomics sample preparation

Each sample was digested in an S-Trap micro spin column (ProtiFi) according to the manufacturer’s instructions. A549 pellets were lysed with RIPA buffer containing protease inhibitors (Pierce). The protein concentration of each sample was measured using a BCA assay (Pierce); 50 µg total protein was used for further analysis. Proteins in each sample were reduced by adding 4.5 mM (final concentration) dithiothreitol and incubated for 15 min at 55°C. Proteins were then alkylated by the addition of 10 mM (final concentration) iodoacetamide for 10 min at room temperature in the dark. Each sample was then acidified by adding phosphoric acid to a 2.5% final concentration. The samples were mixed with 6× volumes of binding buffer (90% methanol; 100 mM TEAB) and loaded onto the S-trap filter and centrifuged at 4,000 × g for 30 s. The samples were washed three times with wash solution (90% methanol; 100 mM TEAB) before digestion with 5 µg trypsin (Promega) (1/10, wt/wt) overnight at 37°C.

The S-trap filters containing digested peptides were then rehydrated for 30 min at room temperature with 40 µL of elution buffer 1 (50 mM TEAB in water). Subsequent elution steps of 40 µL of elution buffer 2 (0.2% trifluoroacetic acid [TFA] in water) followed by 40 µL elution buffer 2 (50% acetonitrile [ACN] in water) completed the peptide elution. The peptide eluates were pooled and de-salted with reverse-phase C18 OMIX tips (Pierce), all according to the manufacturer’s specifications before proceeding to LC-MS/MS analysis.

### LC-MS/MS and data analysis

Peptides were analyzed by LC-MS/MS using a nanoElute coupled to a timsTOF Pro2 Mass Spectrometer (Bruker Daltonics). Samples were loaded on a capillary C18 column (25 cm length, 75 µm inner diameter, 1.6 µm particle size, and 120 Å pore size; IonOpticks). The flow rate was kept at 300 nL/min. Solvent A was 0.1% FA in water, and solvent B was 0.1% FA in ACN. The peptides were separated on a 54.5-min analytical gradient from 2% to 35% of Solvent B for a total of a 60-min run time. The timsTOF Pro2 was operated in the PASEF mode. MS and MS/MS spectra were acquired from 100 to 1,700 m/z. The inverse reduced ion mobility 1/K0 was set to 0.60-1.60 V·s/cm2 over a ramp time of 100 ms. Data-dependent acquisition was performed using 10 PASEF MS/MS scans per cycle with a near 100% duty cycle.

Data analysis was performed with Fragpipe (version 18.0) using the MSFragger (version 3.6) search engine configured with Philosopher (version 4.6.0) and IonQuant (version 1.8.9) with default search settings for LFQ-MBR (match-between-runs) workflow including a false discovery rate set at 1% on both the peptide and protein level. Spectra were compared against Homo sapiens (UniProt ID UP000005640) protein database containing 81,837 human sequences as well as proteins from the H1N1pdm protein database (UniProt ID UP000207659) containing 10 viral protein sequences. For both searches, a mass tolerance for precursor ions was set at a range of −20 ppm to 20 ppm with a mass tolerance for fragment ions at 20 ppm. A maximum of two missed cleavages was set for the shotgun searches. Carbamidomethylation of cysteine residues was set as a fixed modification while variable modifications were set to oxidation of methionine and acetylation of protein N-termini. Only proteins with at least one unique or razor peptide were retained in the shotgun search. Further analysis was performed using the Perseus software v.1.6.2.1 on the proteins to identify significant differences between different genotypes and conditions. Proteins were considered quantified if identified in a minimum of three replicates across all groups, missing values were imputed using default setting in Perseus. Differentially expressed proteins displayed in Figure 4 were selected using 2-way Anova with a p-value cutoff < 0.05 across all comparisons (genotypes, infection and their interaction), meaning significant in all three comparisons. Raw data have been deposited to the ProteomeXchange via PRIDE.

### RNA sequencing and analysis

Stranded RNAseq libraries were constructed using the Kapa mRNA Hyper Prep Kit (Roche). Briefly, the total RNA was quantitated by Qubit (Life Technologies) and assessed for quality with a Fragment Analyzer (Agilent Technologies). Next, polyA+ RNA was selected from 200 ng of total RNA per sample. PolyA+ RNA was fragmented for 4 minutes at 94°C, and then, first-strand cDNA was synthesized with a random hexamer and SuperScript II (Life Technologies). Double-stranded DNA was blunt-ended, 3′-end A-tailed, and ligated to a universal adaptor. The adaptor-ligated double-stranded cDNA was amplified by PCR for 12 cycles with Twist Unique Dual-Index Primers (Twist Bioscience). The final libraries were quantitated on Qubit, and the average size was determined on the Fragment Analyzer and diluted to 5 nM final concentration. The 5 nM dilution was further quantitated by qPCR on a BioRad CFX Connect Real-Time System (Bio-Rad Laboratories Inc). The final stranded RNAseq library pool was sequenced on one lane of an Illumina NovaSeq 6000 S4 flowcell as paired reads with a 150-nt length. The run generated .bcl files which were converted into adaptor-trimmed demultiplexed fastq files using bcl2fastq v2.20 Conversion Software (Illumina, CA). Reads were aligned to the reference genome (GRCh38 (hg38)) and H1N1pdm sequence), and gene-level read counts were generated using Salmon (v1.10.2) for downstream analysis. Differential gene expression analysis was performed using DESeq2, with normalization to the appropriate control group. Genes with statistically significant differential expression were defined as differentially expressed genes (DEGs). Pathway enrichment analysis of DEGs was conducted using the R package clusterProfiler.

### Real-time reverse transcription quantitative PCR (RT-qPCR)

Total cellular RNA was isolated using RNeasy Mini Kit (Qiagen) according to the manufacturer’s manual. One microgram of total RNA was used for the following reverse transcription assays. Reverse transcription (RT) of RNA was performed using ProtoScript II First Strand cDNA Synthesis Kit (New England Biolabs), in accordance with the manufacturer’s manual. Oligo-dT primer was used in RT reaction for detection of mRNAs. Quantification PCR (qPCR) mixtures were prepared according to the user manual of iTaq Universal SYBR Green Supermix (Bio-Rad) and reactions were run in a CFX Opus 96 Real-Time PCR Instrument (Bio-Rad). Segment-specific primers were used for the qPCR analysis.

### Co-immunoprecipitation assay

Transient transfection was performed using PEI, and the DNA: PEI ratio was 1: 3. HEK293T cells that were grown in 6-well plates were co-transfected with 1 μg of TRIM21 expression plasmid together with 1 μg of plasmids encoding gene of interest. Cell lysates were collected 24 hours post transfection. After 24 hours post-transfection, cells were treated with 25 μM MG132 for 6 hours. Immunoprecipitation was performed using either anti-Flag M2 affinity gel (Sigma-Aldrich) or anti-c-Myc magnetic beads (Thermo Fisher Scientific) according to manufacturers’ protocols. The eluates were resuspended in 30 μl of 4× Laemmli Sample Buffer supplemented with β-mercaptoethanol and boiled at 95°C for 10 minutes. The immunoprecipitates were analyzed by Western blot.

### *In vivo* ubiquitination assay

Ubiquitination assay was performed as previously described. Briefly, HEK293T cells cultured in 60 mm dishes were co-transfected with 1 μg of HA-tagged ubiquitin expression plasmids, Myc-TRIM21 and Flag-PR8 NP or Flag-PRKDC using PEI. After 24 hours post-transfection, cells were treated with 25 μM MG132 for 6 hours. The cells were lysed in RIPA lysis buffer (150 mM NaCI, 1% (w/v) TritonX-100, 1 mM EDTA, 0.5% (w/v) sodium deoxycholate, protease inhibitors cocktail (Roche), 50 mM Tris, pH 7.4) containing 100 mM N-ethylmaleimide (NEM) (Sigma-Aldrich). Ubiquitinated protein were precipitated using anti-Flag M2 affinity gel and analyzed by western blot.

### Western blot analysis

Cells were lysed in RIPA lysis buffer and the cell pellets were removed by centrifugation at speed of 14,000 rpm for 30 min at 4 °C. Protein samples were prepared by mixing with 4× Laemmli Sample Buffer (Bio-Rad) supplemented with β-mercaptoethanol (Sigma-Aldrich) and boiled at 95 °C for 10 min. After SDS-PAGE, the proteins were transferred from the gel to polyvinylidene fluoride membranes (Bio-Rad). Membranes were blocked with 5% skim milk in PBST (PBS supplemented with 0.1% Tween-20) for 1 h at room temperature and incubated overnight with primary antibodies diluted in 5% skim milk in PBST at 4°C. Membranes were washed three times, 10 min each with PBST, incubated with secondary antibodies for 1 hour at room temperature. Afterwards, membranes were washed three times, 10 min each with PBST. Positive immunostaining bands on the membranes were visualized using ECL Select Western Blotting Detection Reagent (Cytiva) and scanned by iBright 1500 imaging system (Invitrogen).

### Immunofluorescence assay

One day before the experiment, cells were seeded on coverslips pre-coated with Poly-L-Lysine (Sigma-Aldrich) at a density of 0.3 x 10^6^ cells/well in 12-well plates. Cells were infected by viruses at a MOI of 10. Coverslips with cells were collected and washed twice with PBS, followed by fixation with 4% formaldehyde for 30 minutes at room temperature. The cells were permeabilised with 0.5% Triton, followed by two washes with PBS. The cells were then blocked with 5% normal goat serum for 30 minutes at room temperature. Primary antibodies at their appropriate dilution in PBS was added and incubated for 1 hr at room temperature followed by two washes with PBS. Secondary antibodies diluted in PBS was dispensed and incubated for 30 minutes at room temperature. After three washes, the coverslips were mounted on glass slides with DAPI Fluoromount-G (Southern Biotech) and kept at 4°C until visualization with confocal microscopy.

### Fluorescence in situ hybridization

One day before the experiment, cells were seeded on coverslips in 24-well plates. Cells were infected by viruses at an MOI of 10. Infection was synchronized and secondary infection was blocked using 20 mM NH_4_CI. Coverslips with cells were collected and washed twice with PBS, followed by fixation with 4% formaldehyde for overnight at 4 °C. To visualize viral RNA, RNA fluorescence in situ hybridization (RNA-FISH) through hybridization chain reaction (HCR)^78^ was then performed according to the manufacturer’s instructions with modifications. Cells were washed with PBS thrice, then permeabilized with 0.1% v/v Triton X-100 for 10 minutes at room temperature. After the permeabilization, cells were washed twice with 2× sodium chloride sodium citrate (SSC) buffer and incubated in HCR hybridization buffer at 37 °C for 30 minutes with gentle rocking. Subsequently, cells were incubated with hybridization buffer containing 1.2 pmol of the PB2 and NP vRNA probe at 37°C for overnight with gentle rocking.

The next day, excess probes were removed by washing cells 4 times with probe wash buffer at 37 °C with gentle rocking and were further washed twice with 5× SSC-T buffer (5× SSC with 0.1% Tween-20) for 5 minutes each at room temperature. Cells were then incubated with amplification buffer for 30 min at room temperature with gentle rocking. 18 pmol of B1-Alexa 488 or B3-Alexa 555 amplifier hairpins per sample were snap-cooled by heating at 95°C for 90 seconds and cooled to room temperature in the dark for 30 minutes. Hairpin solution, which was prepared by adding amplifier hairpins to 300 μL amplification buffer per sample, was added to samples and incubated for 1 hour at room temperature in the dark with gentle rocking. Excess hairpins were removed by washing samples 5 times with 5× SSC-T buffer. For the first wash, samples were incubated in 5× SSC-T buffer supplemented with 1 μg/mL DAPI (Invitrogen) for 20 minutes at room temperature with gentle rocking. All subsequent washes were performed for 5 min at room temperature with gentle rocking. After the last wash, samples were mounted on glass slides with ProLong Diamond Antifade Mountant (Invitrogen). Cells were visualized using a LSM880 microscope system (Zeiss).

### Positive selection analysis

TRIM21 sequences were obtained from NCBI and aligned using MUSCLE as implemented in MEGA 11^79^. Gaps in alignments were trimmed using Gblocks v1.0^80^ with default parameters, except gap positions was allowed to half. Species trees were inferred using TimeTree. Signatures of evolutionary selection were assessed using the CodeML program in the PAML software package PAML^81^. Analyses were performed using site models with the F3×4 codon frequency model. Likelihood ratio tests were used to compare Model 8 (beta and omega-allowing for positive selection) and Model 7 (beta-no positive selection). Statistical significance was determined using a chi-squared test on twice the difference in log-likelihood values (2ΔlnL) between models, with two degrees of freedom. To ensure convergence of likelihood estimates, each analysis was performed using two different initial ω (dN/dS) values (0.4 and 1.5). Accession numbers for input sequences can be found in **Table S2**.

### Quantification and statistical analysis

All the statistical analyses have been performed using Prism 9 Graph Pad Software. Two-tailed Student’s unpaired t-test was performed to compare between two populations of data (e.g., control and sample) whereas two-way ANOVA was applied for multi sample comparisons. All data generated were from independent biological replicates where n ≥ 3, each measured in technical duplicates or triplicates. Results have been presented as means ± standard deviation (SD) or standard error mean (SEM).

## Data availability

Mass spectrometry proteomics data have been submitted to the ProteomeXchange Consortium via the PRIDE repository. The dataset identifier will be made available upon publication.

