## Supplementary material for "TRIM21 is a molecular rheostat for influenza A virus replication": SI figure

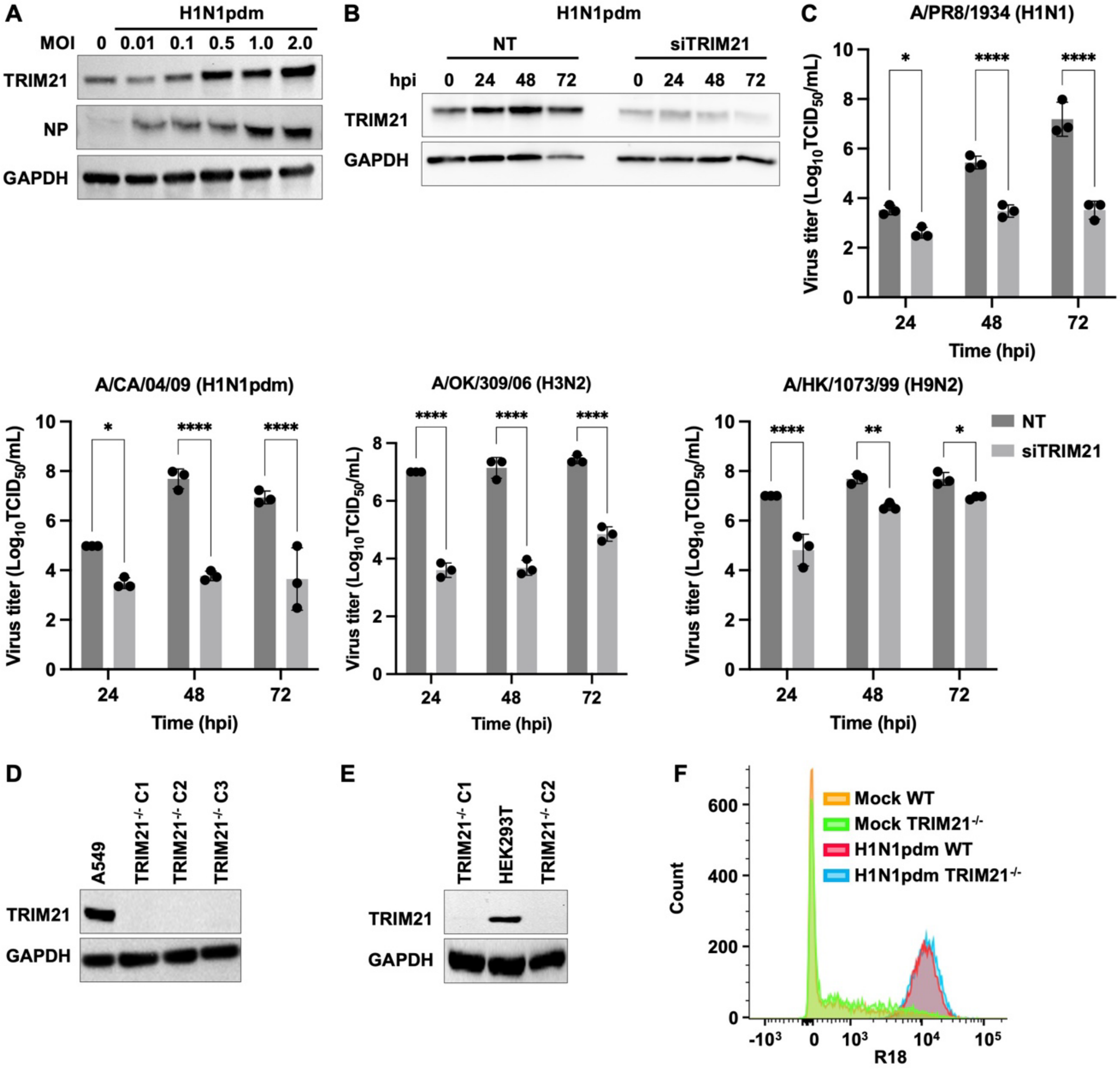


**Figure S1. TRIM21 deficiency inhibits viral production. (A)** A549 cells were infected with H1N1pdm at varying MOI (0.01, 0.1, 0.5, 1 and 2). Lysates prepared from infected cells were immunoblotted with antibodies against TRIM21 and NP. GAPDH was used as a loading control. **(B)** TRIM21 was depleted in A549 cells by siRNA-mediated knockdown. Non-targeting control (NT) and TRIM21-depleted (siTRIM21) cells were subsequently infected with H1N1pdm (MOI = 0.01). Knockdown efficiency was verified by immunoblotting using anti-TRIM21 antibody. **(C)** Viral titers in NT and siTRIM21 A549 cells infected with H1N1, H1N1pdm, H3N2, or H9N2 viruses (MOI = 0.01) at the indicated time points were measured by $TCID_{50/mL}$. Data are shown as means of n = 3 ± standard deviations (SD). Two-way ANOVA was used to analyze data (*$p < 0.05$, **$p < 0.01$, ****$p < 0.0001$). **(D and E)** Western blot validating CRISPR/Cas9-mediated TRIM21 deletion ($TRIM21^{-/-}$) in **(D)** A549 and **(E)** HEK293T cells. **(F)** Quantification of R18-positive population in mock and infected A549 cells, as measured by flow cytometry.

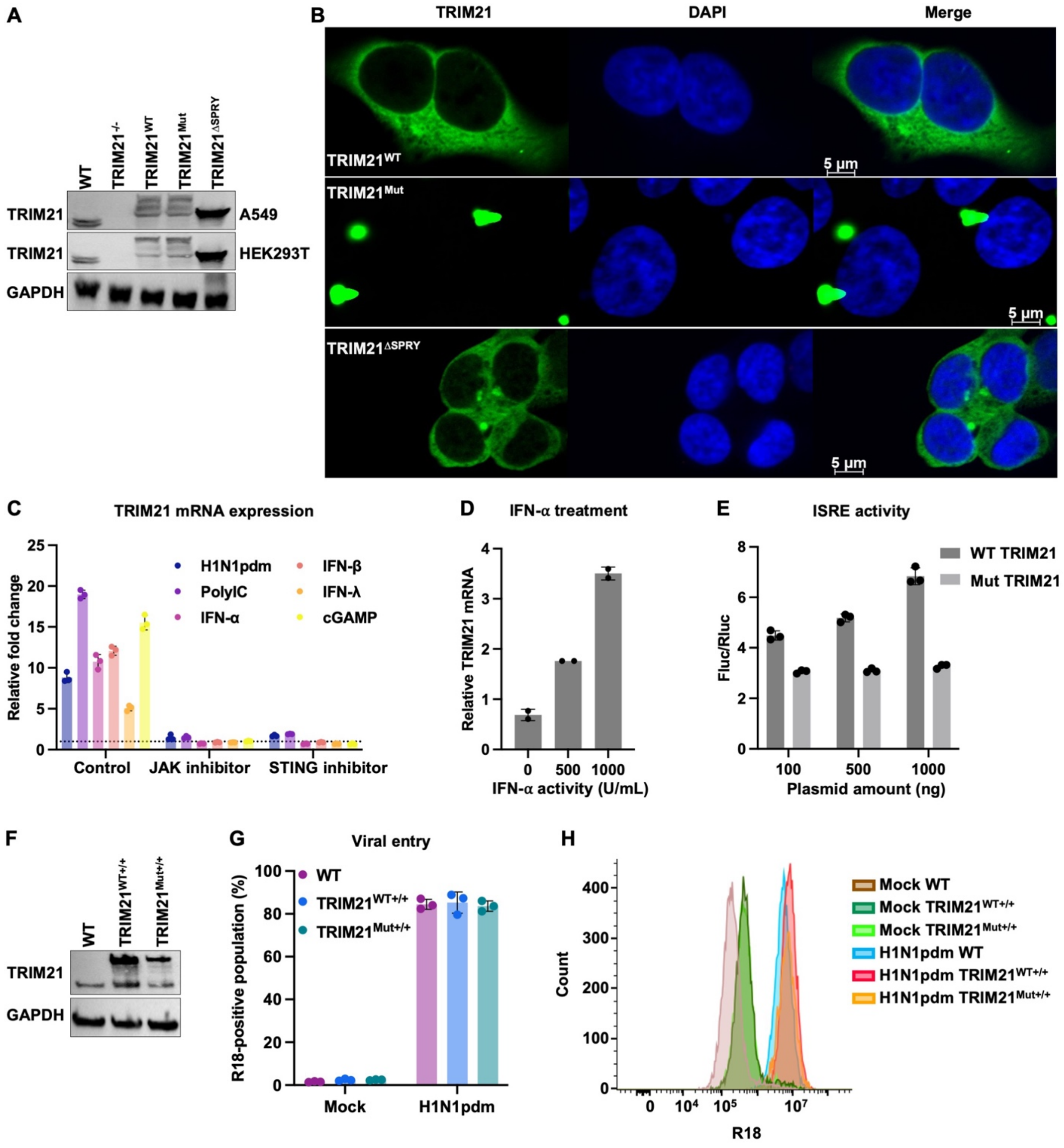


**Figure S2. TRIM21 suppresses IAV replication by coupling NP degradation to innate immune amplification. (A)** Immunoblot validation of A549 and HEK293T cells reconstituted with WT TRIM21 (TRIM21$^{WT}$), a catalytically inactive mutant of TRIM21 (TRIM21$^{Mut}$) or a SPRY domain-deficient TRIM21 (TRIM21$^{\Delta SPRY}$). TRIM21 constructs were GFP-tagged and therefore migrated at a higher molecular weight than endogenous TRIM21 (55 kDa). **(B)** Immunofluorescence imaging of TRIM21 subcellular localization in the indicated reconstituted A549 cell lines. **(C)** A549 cells were treated with the indicated innate immune stimuli in the presence or absence of pathway-specific inhibitors. Total RNA was isolated, and TRIM21 mRNA levels were quantified by real-time PCR. Data are shown as means of n = 3 ± standard deviations (SD). The dotted line indicates fold change of 1. **(D)** A549 cells were treated with increasing concentrations of IFN-α, followed by analysis of TRIM21 mRNA expression by real-time PCR.

Data are shown as means of n = 2 ± standard deviations (SD). **(E)** HEK293T cells were transiently transfected with increasing amounts of plasmids expressing WT Myc-TRIM21 or the catalytically inactive mutant Myc-TRIM21, together with an ISRE-driven firefly luciferase reporter and a Renilla luciferase control. At 24 hours post-transfection, cells were treated with IFN-α for an additional 24 hours prior to measurement of luciferase activities. Data are shown as means of n = 3 ± standard deviations (SD). **(F)** Immunoblot validation of A549 cells stably overexpressing WT TRIM21(TRIM21$^{WT+/+}$) or the catalytically inactive mutant (TRIM21$^{Mut+/+}$). TRIM21 constructs were GFP-tagged and therefore migrated at a higher molecular weight than endogenous TRIM21 (55 kDa). **(G and H)** Viral entry was assessed by flow cytometric analysis of R18-positive populations at 1 hpi. Data are shown as means of n = 3 ± standard deviations (SD).

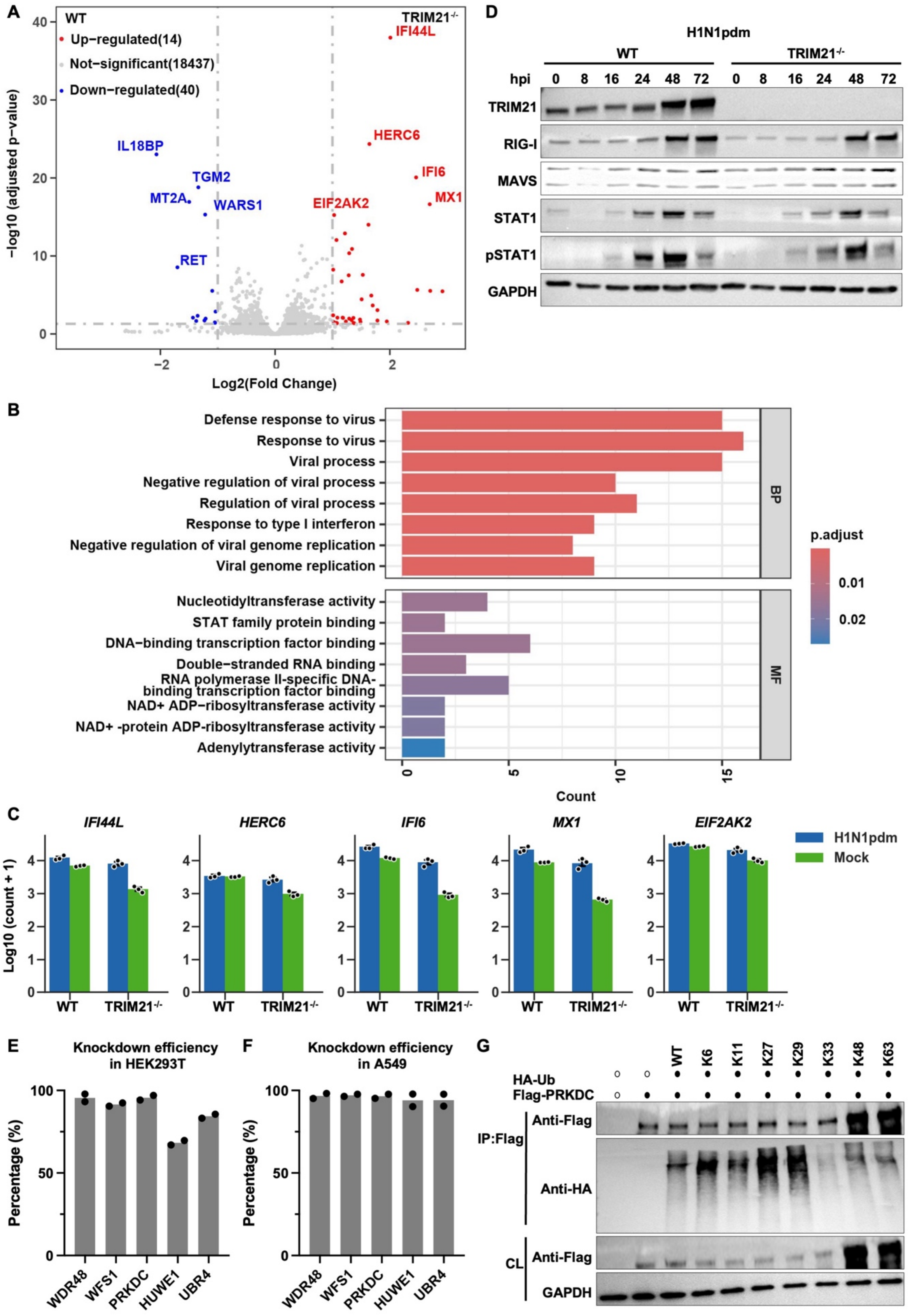

A
WT
Up−regulated(14)
Not−significant(18437)
Down−regulated(40)
TRIM21-/-
IFI44L
HERC6
IFI6
MX1
EIF2AK2
IL18BP
TGM2
MT2A
WARS1
RET
−log10 (adjusted p−value)
Log2(Fold Change)
B
Defense response to virus
Response to virus
Viral process
Negative regulation of viral process
Regulation of viral process
Response to type I interferon
Negative regulation of viral genome replication
Viral genome replication
Nucleotidyltransferase activity
STAT family protein binding
DNA−binding transcription factor binding
Double−stranded RNA binding
RNA polymerase II-specific DNA-binding transcription factor binding
NAD+ ADP−ribosyltransferase activity
NAD+ -protein ADP-ribosyltransferase activity
Adenylyltransferase activity
BP
MF
p.adjust
Count
C
IFI44L
HERC6
IFI6
MX1
EIF2AK2
H1N1pdm
Mock
Log10 (count + 1)
WT
TRIM21-/-
D
H1N1pdm
WT
TRIM21-/-
hpi 0 8 16 24 48 72 0 8 16 24 48 72
TRIM21
RIG-I
MAVS
STAT1
pSTAT1
GAPDH
E
Knockdown efficiency in HEK293T
F
Knockdown efficiency in A549
Percentage (%)
WDR48
WFS1
PRKDC
HUWE1
UBR4
G
WT K6 K11 K27 K29 K33 K48 K63
HA-Ub
Flag-PRKDC
IP:Flag
Anti-Flag
Anti-HA
CL
Anti-Flag
GAPDH

**Figure S3. Loss of TRIM21 reveals a PRKDC-dependent compensatory antiviral program.** **(A)** Volcano plot showing differentially expressed genes (DEGs) between H1N1pdm-infected WT and TRIM21$^{-/-}$ A549 cells (MOI 0.01, 48hpi). The x-axis indicates $\log_2$ fold change and the y-axis shows $-\log_{10}$ adjusted *p*-value. DEGs were defined using an adjusted *p*-value cutoff of 0.05 and an absolute $\log_2$ fold change cutoff of 1. Colors indicate upregulated (red) or downregulated (blue) genes. **(B)** Gene Ontology analysis for upregulated DEG in biological process (BP) and molecular function (MF) in H1N1pdm-infected TRIM21$^{-/-}$ A549 cells. The number on the x-axis indicates the enriched count of DEG. **(C)** Transcriptomic raw count of ISGs in mock- and H1N1pdm-infected WT and TRIM21$^{-/-}$ A549 cells (MOI 0.01, 48hpi). **(D)** WT and TRIM21$^{-/-}$ A549 cells were infected with H1M1pdm (MOI = 0.01), and cell lysates were immunoblotted for TRIM21, RIG-I, MAVS, STAT1, and phosphorylated (pSTAT1). **(E and F)** siRNA-mediated knockdown efficiency of the indicated targets in **(E)** HEK293T and **(F)** A549 cells measured by real-time PCR. **(G)** HEK293T cells were transiently co-transfected with Flag-PRKDC and HA-ubiquitin (WT versus linkage-specific constructs as indicated). Cells were treated with MG132, followed by immunoprecipitation using anti-Flag M2 affinity beads. IP and CL were then analyzed by immunoblotting. Black and white dots indicate the presence or absence of plasmid transfection, respectively.

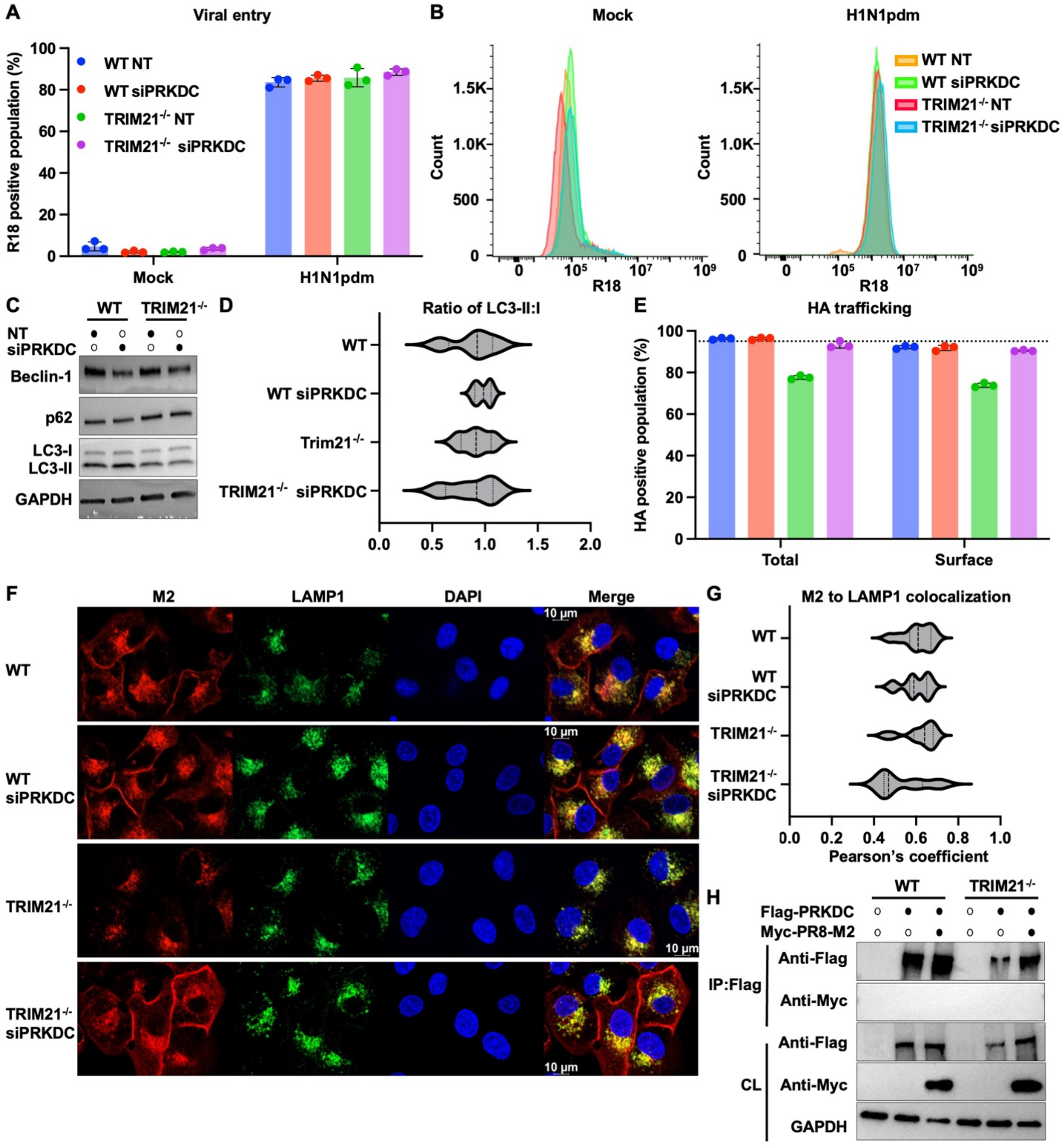


**Figure S4. PRKDC-mediated restriction is independent of viral entry and autophagic degradation of viral proteins. (A and B)** Viral entry was assessed by flow cytometric analysis of R18-positive populations at 1 hpi. Data are shown as means of n = 3 ± standard deviations (SD). **(C)** WT and TRIM21$^{-/-}$ A549 cells were transfected with NT or PRKDC-targeting siRNA and subsequently infected with H1N1pdm (MOI = 0.01). Lysates were immunoblotted Beclin-1, p62, and LC3. The cell lines are indicated by black dots. **(D)** Ratio of LC3-II to LC3-I were quantified by densitometry. Each violin represents *n*= 4 independent samples. **(E)** WT and TRIM21$^{-/-}$ A549 cells were transfected with NT or PRKDC-targeting siRNA and subsequently infected with H1N1pdm (MOI = 10). Total and surface expression of viral HA protein were quantified by flow cytometry. The dotted line indicates the mean HA expression in

NT-treated WT A549 cells. Data are shown as means of n = 3 ± SD. **(F)** Immunofluorescence imaging of viral M2 and the lysosomal marker LAMP1 were performed in indicated H1N1pdm-infected A549 cells (MOI = 10). (**G)** Colocalization coefficient between M2 and LAMP1 in the indicated H1N1pdm infected A549 cells was quantified. Each violin represents *n* = 11 images, except for TRIM21$^{-/-}$ siPRKDC (*n* = 10). **(H)** HEK293T cells were transiently co-transfected with Flag-PRKDC and Myc-PR8-M2. Lysates were immunoprecipitated with anti-Flag M2 affinity beads and analyzed by immunoblotting. Black and white dots indicate the presence or absence of plasmid transfection, respectively.

**A** Positive selection analysis

| Mammalian order | | Starting omega 0.4 | Starting omega 1.5 |
|---|---|---|---|
| Primates | Full | 0.09 | 0.09 |
| | SPRY | 0.21 | 0.21 |
| Rodents | Full | 0.02 | 0.21 |
| | SPRY | 0.31 | 0.31 |
| Carnivores | Full | 0.19 | 0.19 |
| | SPRY | 0.54 | 0.19 |
| Even-toed ungulates | Full | 0.24 | 0.24 |
| | SPRY | 0.99 | 0.99 |

**B** Multiple sequence alignment of TRIM21 RING domain

```
Human       CPICLDPFVEPVSIECGHSFCQECI
Chimpanzee  CPICLDPFVEPVSIECGHSFCQECI
Macaque     CPICLDPFVEPVSIECGHSFCQECI
Mouse       CSICLDPMVEPMSIECGHCFCKECI
Rat         CSICLEPMVEPMSIECGHCFCKECI
Ferret      CPICLDPMVEPMSIDCGHNFCHECI
Hamster     CSICLDPMVAPMSIECGHCFCKECI
Pig         CSICLDPMVEPVSIECGHSFCQECI
Seal        CPICLDPMVEPMSIDCGHSFCHECI
```

**C** Multiple sequence alignment of TRIM21 SPRY domain

```
Human       ELRSVCHVPGLKKMLRTCAVHITLDPDTANPWLILSEDRRQVRLGDTQQSIPGNEERFDSYPMVLGAQHFHSG
Chimpanzee  ELRSVCHVPGLKKMLRTCAVHITLDPDTANPWLILSEDRRQVRLGDTQQSIPGNEERFDSYPMVLGAQHFHSG
Macaque     ELRSVCHVPGLKKMLRTCAVHITLDPDTANPWLILSEDRRQVRLGDTQQSIPGNEERFDSYPMVLGAQHFHSG
Mouse       DLTSTCPVPGRKKMLRTCWVHITLDRNTANSWLIISKDRRQVRMGDTHQNVSDNEERFSNYPMVLGAQRFSSG
Rat         DLTSTCHVPGRKKMLRTCWVHITLDRDTANSCLIISKDRRQVRMGDTYQNVSENAERFNNYPMVLGAQRFSSG
Ferret      DLRNVCLVPGLKRMLRTCGVHITLDPHTANPWLILSEDRRQVRLGDRRQEVPENEDRFDHYPMVLGAQCFDSG
Hamster     DLTNACHVPGRKKMLRTCWVHITLDRNTANPWLIIPKDRRQVRIGDSHQNVSENDERFNNYPVVLGAQRFTSG
Pig         DLSILCYVPGIKRMLRSYAVHITLDPLTASPWLVLSENRRQVRLGETRQEVPENEERFDTYPMVLGAQHFNSG
Seal        DLRSVCLVPGLKRMLRTCGVHITLDPHTANPWLILSKDQRQVRLGDSQQKVPENEDRFDSYPMVLGAQCFDSG

Human       KHYWEVDVTGKEAWDLGVCRDSVRRKGHFLLSSKSGFWTIWLWNKQKYEAGTYPQTPLHLQVPPCQVGIFLDY
Chimpanzee  KHYWEVDVTGKEAWDLGVCRDSVRRKGHFLLSSKSGFWTIWLWNKQKYEAGTYPQTPLHLQVPPCQVGIFLDY
Macaque     KHYWEVDVTGKEAWDLGVCRDSVRRKGHFLLSSKSGFWTIWLWNKQKYEAGTYPQTPLHLQVPPCQIGIFLDY
Mouse       KMYWEVDVTQKEAWDLGVCRDSVQRKGQFSLSPENGFWTIWLW-QDSYEAGTSPQTTLHIQVPPCQIGIFVDY
Rat         KMYWEVDVTQKQAWDLGVCRDSVQRKGHFSLSPENGFWTIWLWQKDNYEAGTSPQTTLHIQVPPCQIGIFVDY
Ferret      QVYWEVDVTGKEAWDLGVCRDSVQRKGQFSLSPETGFWTIWLWNKQKYEAGTCIQTPLHLQVPPRQVGIFLDY
Hamster     KMYWEVDVTRKEAWELGICRESVQRKGQFLFSPENGFWIIWLWNKDIYEAGTNPQTTLHLQVPPCQVGIFVDY
Pig         KHYWEVDVAGKEAWDLGVCRDSVKRKGHFLLNPSNGFWTIWLWNKQKYEAGTCTQTPLYPQVPPSRVGIFLDY
Seal        KVYWEVDVTGKEAWDLGVCRDSVQRKGHFLLSPENGFWTIWLWNKQKYEAGTCPQTPLHLQVPPRQVGIFLDC

Human       EAGMVSFYNITDHGSLIYSFSECAFTGPLRPFFSPGFNDGGKNTAPLTLCPL
Chimpanzee  EAGMVSFYNITDHGSLIYSFSECAFTGPLRPFFSPGFNDGGKNTAPLTLCPL
Macaque     EAGMVSFYNITDHGSLIYSFSECAFTGPLRPFFSPGFNDGGRNTAPLTLCPL
Mouse       EAGVVSFYNITDHGSLIYTFSECVFAGPLRPFFNVGFNYSGGNAAPLKLCPL
Rat         EAGIVSFYNITDHGSLIYTFSECAFIGPLRPFFSVGFNDDGGNSAPLKLCPL
Ferret      EARIVSFYNITDHGSLIYTFSECAFAEPLRPFFSTGFNDGGRNAAPLVLCPL
Hamster     EAGIVSFYNITDHGSLIYTFSQCTFAGPLRPFFSVGFNDDGGNAAPLKLCPL
Pig         DASTVSFYNISDHGSLIYSFSECAFAGPVRPFFNPGFNDGGRNAPPLTLCPL
Seal        EARIVSFYNVTDHGSLIYTFSECAFAGPLRPFFSTGFNDEGRNGAPLVLCPL
```

**Figure S5. Comparative evolutionary analysis of TRIM21 across mammals. (A)** Positive selection analysis of full-length and SPRY domain of TRIM21 in the indicated mammalian order. Analyses were performed using orthologous TRIM21 sequences listed in Table S2. **(B and C)** Multiple sequence alignment of TRIM21 from representative species highlighting conservation within the **(B)** RING domain and **(C)** SPRY domain. Catalytic residues and residues implicated in NP binding are marked with red.
